# Spermatogenic context controls outcomes of engineered sex distortion in malaria mosquitoes

**DOI:** 10.64898/2026.03.29.715086

**Authors:** Lee Benjamin Lamdan, Sigal Popovsky-Sarid, Ebrima SM Kolley, Arad Sarig, Daniella An Haber, Elad Shmuel Yonah, Eric Marois, Leonidas-Romanos Davranoglou, Yael Arien, Philippos Aris Papathanos

## Abstract

Sex-ratio distortion systems are promising genetic tools for mosquito population control. Two strategies have been proposed: prezygotic elimination of X-bearing sperm by X-shredding, which can drive invasive Y-chromosome transmission when sex distorters are Y-linked, and postzygotic daughter killing through disruption of X-linked haploinsufficient genes, a self-limiting approach known as X-poisoning. Previous attempts to implement X-poisoning in the malaria mosquito *Anopheles gambiae* unexpectedly produced prezygotic distortion, with sex bias arising from loss of X-bearing sperm rather than daughter lethality.

Here we use a split CRISPR-Cas9 system to systematically compare sex-ratio distortion outcomes across germline Cas9 drivers and X-linked target genes. Meiotic X-chromosome targeting induced preferential Y-chromosome transmission regardless of target identity, function, or number of sgRNA target sites. In contrast, shifting Cas9 expression to earlier spermatogenic stages altered outcomes dramatically: targeting X-linked ribosomal protein genes caused severe developmental or reproductive toxicity, whereas targeting the haplolethal muscle gene *wupA* produced daughter-specific post-embryonic lethality, with the majority of surviving females emerging flightless. Tracking offspring using a Y-linked fluorescent marker confirmed that sex chromosome segregation remained unbiased, with female mortality accumulating progressively from the first larval instar, reaching near-complete lethality by adulthood.

These results demonstrate that the timing of Cas9 expression during spermatogenesis, rather than target gene identity alone, determines the outcome of X-chromosome targeting in malaria mosquitoes, and establish the conditions required for genuine X-poisoning. Identification of *wupA* as an effective X-poisoning target provides a solid foundation for the future development of self-limiting Y-linked sex-ratio distortion systems for malaria vector control.

## Introduction

In 1967, Hamilton proposed that a Y-linked element capable of biasing gamete production in its own favor could form the basis of a new type of mosquito genetic control strategy. In theory, such an “extraordinary” Y chromosome could spread through a population, progressively skewing the sex ratio toward males and ultimately causing population collapse due to the lack of females (Hamilton, 1967). Hamilton’s framework drew on earlier cytological observations of naturally occurring sex-ratio distorters discovered in *Aedes aegypti* mosquitoes, where fragmentation of the X chromosome during male meiosis was associated with an excess of male offspring (Hickey and Craig, 1966). Burt later proposed that such a sex-ratio distorter (SRD) could be engineered by harnessing site-specific endonucleases to selectively target X chromosomes during spermatogenesis (Burt, 2003).

In the malaria vector *Anopheles gambiae*, proof-of-concept SRDs were first developed using the meganuclease I-*Ppo*I and later CRISPR-Cas9 to target the repetitive X-linked 28S rDNA array during spermatogenesis (Windbichler, Papathanos and Crisanti, 2008; Galizi *et al*., 2014, 2016). Cleavage of the rDNA array in developing sperm interferes with the transmission of targeted X chromosomes and results in preferential inheritance of Y-bearing gametes, a mechanism we called X-shredding. X-shredding transgenic males sire a strong excess of male offspring (>95%), while largely preserving reproductive output (Galizi *et al*., 2014, 2016). More recently, an alternative strategy termed X-poisoning was proposed, in which Cas9 targets single-copy X-linked haploinsufficient (HI) genes (Fasulo *et al*., 2020). In this system, cleavage of the paternal X chromosome generates loss-of-function alleles that are transmitted through otherwise functional sperm, but confer lethality to daughters inheriting the disrupted paternal X chromosome. As a result, sex-ratio distortion arises through postzygotic lethality of daughters rather than prezygotic bias of sex chromosome transmission.

When linked to the Y chromosome, these two SRD mechanisms have fundamentally different population dynamics (Burt and Deredec, 2018). In a Y-linked X-shredder, linkage of the SRD to the Y chromosome couples preferential production of Y-bearing sperm with its own transmission, such that the over-represented male offspring all inherit the SRD allele. This mutually beneficial arrangement enables the SRD to spread through the population in an invasive manner. By contrast, X-poisoning does not alter the ratio of X- and Y-bearing sperm. Instead, sex-ratio distortion arises through the selective loss of daughters during development. As a result, the frequency of the SRD allele remains tied to the initial release frequency, making Y-linked X-poisoning, alternatively called Y-linked editors (YLEs), intrinsically self-limiting (Burt and Deredec, 2018). Nevertheless, linkage to the Y chromosome insulates the SRD from negative selection in females, allowing greater persistence than autosomal SRDs, which are rapidly eliminated due to the fitness costs imposed on daughters. This self-limiting character may be advantageous in contexts where self-sustaining spread is undesirable.

Experimental work in *D. melanogaster* demonstrated that X-poisoning can be engineered using CRISPR-Cas9, and that target copy number is a key determinant of the distortion mechanism (Fasulo *et al*., 2020). Fasulo et al. showed that meiotic β*2-tubulin*-driven Cas9 targeting of the repetitive multi-copy *Muc14a* locus produced prezygotic X-chromosome elimination. By contrast, applying X-poisoning through the targeting of single-copy X-linked ribosomal protein genes produced daughter-specific postzygotic lethality, with sex ratios exceeding 93% males. These results confirmed the prediction that copy number determines whether Cas9 activity results in X-chromosome elimination or in the transmission of disrupted alleles to daughters who then die in postzygotic stages. Lawler et al. (2024) subsequently extended X-poisoning to the single-copy, cytoskeletal muscle gene *wings up A* (*wupA*), which produced strong female-specific lethality late in embryogenesis when *wupA* was targeted during spermatogenesis and transmitted to daughters (Lawler *et al*., 2024). Together, these studies established that single-copy haploinsufficient X-linked genes are effective targets for postzygotic sex-ratio distortion through selective loss of daughters.

Based on this model, we previously attempted to engineer X-poisoning in *An. gambiae* using transgenes in which Cas9 and sgRNA components were encoded together in a single construct and inserted into the genome via *piggyBac*-mediated transposition (Haber *et al*., 2024). Unexpectedly, we found that even when targeting single-copy X-linked ribosomal protein genes, the outcome was not postzygotic daughter lethality but rather the elimination of X-bearing sperm, effectively mimicking X-shredding, a mechanism previously thought to require targeting of repetitive sequences. At the time, these experiments used only the meiotic β*2-tubulin* (β*2t*) promoter, as it had proven effective for X-poisoning in *D. melanogaster* (Fasulo *et al*., 2020) and also for X-shredding in other insect species (Meccariello *et al*., 2021). However, because transgenes were integrated at random genomic positions, it was difficult to directly compare outcomes across different lines, sgRNA designs, or target genes. Together, these constraints left open the question of whether the failure to achieve X-poisoning in *An. gambiae* reflected the Cas9 driver, the target, or some combination of both.

To address these limitations, here we re-established the X-poisoning system in *An. gambiae* using a split CRISPR-Cas9 design in which Cas9 and sgRNA components are encoded on separate transgenes and inserted at well-characterized autosomal attP sites. This approach allowed us to independently compare two germline Cas9 drivers, the meiotic β*2t* promoter and the pre-meiotic *zero population growth* (*zpg*) promoter, and three X-linked target genes, including two previously best-performing ribosomal protein gene targets and the *An. gambiae wupA* ortholog (**Figure 1A**), while keeping transgene insertion sites constant across comparisons. We show that the developmental stage at which Cas9 is active during spermatogenesis determines the outcome of X-linked targeting in *An. gambiae*. When Cas9 is expressed in late primary spermatocytes undergoing meiosis by the β*2t* promoter (Santel *et al*., 2000; Catteruccia, Benton and Crisanti, 2005), X-chromosome cleavage leads to elimination of X-bearing sperm regardless of target identity. In contrast, earlier expression of Cas9 by the *zpg* promoter in germline stem cells and mitotic spermatogonia prior to meiosis (Tazuke *et al*., 2002; Hammond *et al*., 2021), enables transmission of disrupted X-linked alleles and produces genuine postzygotic daughter lethality. These findings explain why previous attempts to engineer X-poisoning in mosquitoes failed and establish conditions under which this strategy can be achieved.

**Figure 1.**
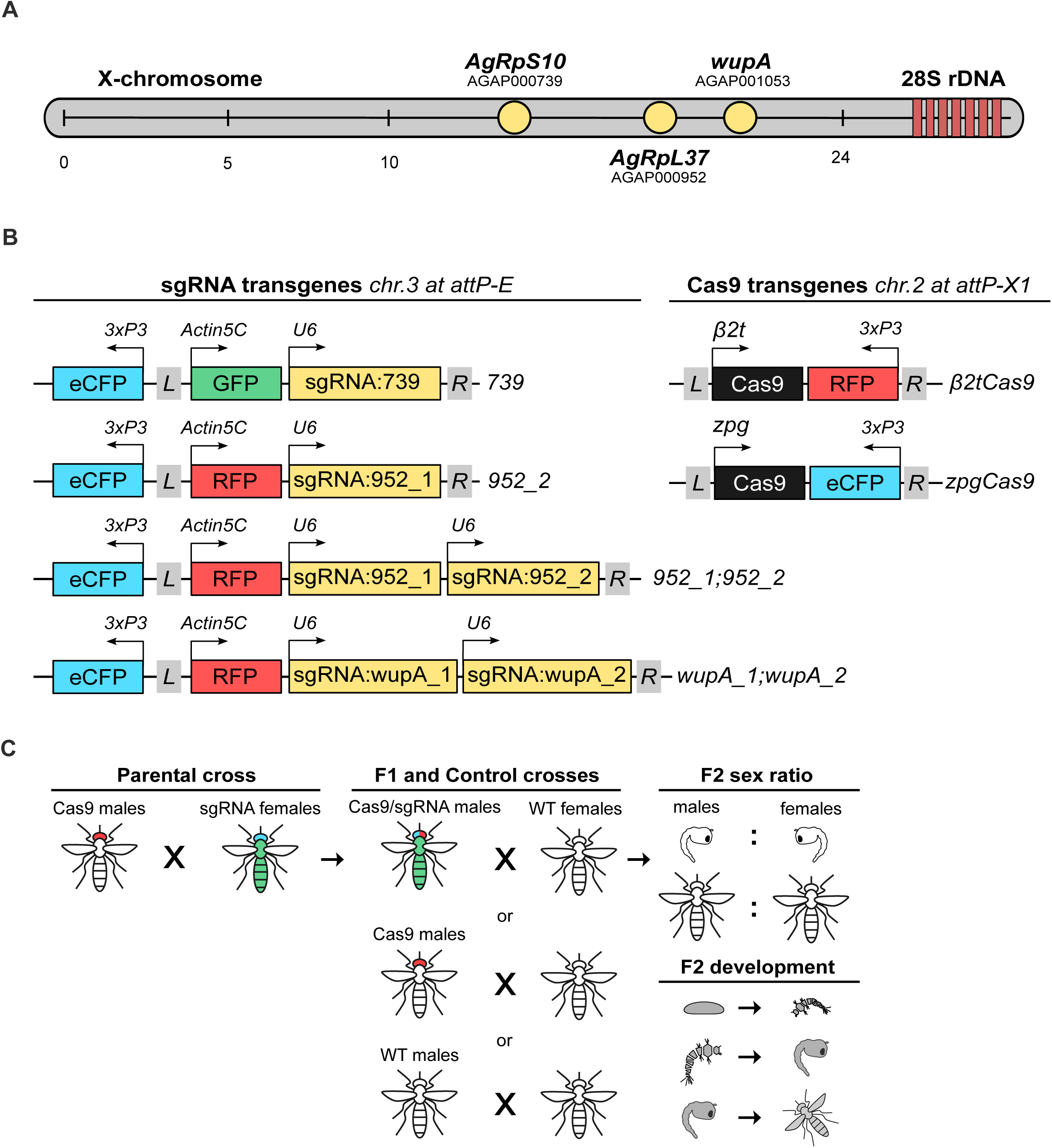
Experimental design of the split CRISPR-Cas9 sex-ratio distortion system. (**A**) Schematic of the X chromosome of *Anopheles gambiae* showing the positions of the three X-linked target genes (*AgRpS10*, *AgRpL37*, and *wupA*) and the 28S rDNA cluster used for X-shredding. (**B**) Transgenic constructs generated. Left: sgRNA transgenes integrated at the autosomal attP-E docking site on chromosome 3, carrying either eGFP or RFP markers driven by the *An. gambiae Actin5C* promoter, and one or two sgRNA expression cassettes driven by the *U6* promoter. Right: Cas9 transgenes integrated at the attP-X1 docking site on chromosome 2, expressing Cas9 under the meiotic β*2-tubulin* promoter (β2Cas9, with 3xP3:RFP marker) or the pre-meiotic *zpg* promoter (zpgCas9, with 3xP3:eCFP marker). (**C**) Crossing scheme. Parental crosses between homozygous Cas9 males and homozygous sgRNA females produce transheterozygous F1 males. These are then backcrossed to wild-type females to generate F2 offspring whose sex ratio and stage-specific survival are scored across development.

## Results

### A split CRISPR-Cas9 system to compare germline Cas9 drivers and X-chromosome targets

To enable a controlled comparison of X-linked sex-ratio distortion phenotypes across different Cas9 drivers, we generated a split (two-component) transgenic CRISPR-Cas9 system in which Cas9 and sgRNA components are encoded on separate transgenes and are integrated at defined attP docking sites. A β*2-tubulin*-driven Cas9 (hereafter β*2*Cas9) construct, carrying a 3xP3:RFP transformation marker, was integrated into the markerless attP-X1 docking site on chromosome 2L (Volohonsky *et al*., 2015) (**Figure 1B**). In parallel, we generated a series of sgRNA lines targeting X-linked ribosomal protein genes, using our best-in-class sgRNAs against *AgRpS10* (AGAP000739) and *AgRpL37* (AGAP000952) from Haber et al., 2024, implemented as single- or double-sgRNA designs. sgRNA constructs carried either Actin5C:eGFP or Actin5C:RFP fluorescent markers and were integrated into the attP-E docking site on chromosome 3R (Meredith *et al*., 2011) (**Figure 1B**).

To assess sex-ratio distortion, homozygous β*2*Cas9 males were crossed to each sgRNA line to generate transheterozygous F1 males. These were then backcrossed to wild-type females, and F2 phenotypes were scored from egg to adult stages (**Figure 1C**). All β*2*Cas9/sgRNA combinations produced significant male bias relative to controls at both the pupal and adult stages (one-way ANOVA, *p* < 0.0001; **Figure 2A**). Targeting *AgRpL37* using the 952_2 and 952_1;952_2 sgRNAs yielded the strongest distortion, with male frequencies of 96-97%, while targeting *AgRpS10* (sgRNA-739) resulted in ∼86% males. There were no substantial differences between sex ratios at the pupal and adult stages, indicating no detectable sex-specific mortality during late development. Offspring from control crosses exhibited near-equal sex ratios in both developmental stages (**Figure 2A**). To determine whether pre- or postzygotic mechanisms underlie the observed sex bias, we quantified survival across developmental stages, reasoning that postzygotic lethality would manifest as stage-specific excess mortality in the experimental crosses relative to controls (**Figure 2B**). Egg hatching rates were uniformly high across all genotypes indicating no detectable impact on embryonic viability. Larval-to-pupal survival was modestly reduced in offspring from the cross targeting *AgRpL37* with the dual-sgRNA (952_1;952_2) (Welch’s one-way ANOVA, *p* < 0.0001), but this reduction did not scale with the magnitude of sex distortion observed at later stages. Pupal-to-adult survival was comparable across all β*2*Cas9/sgRNA combinations and controls (**Figure 2B**). These results were consistent with our previous findings using combined β*2*Cas9/sgRNA constructs (Haber *et al*., 2024) and confirmed the absence of differential developmental lethality between the sexes, suggesting that the observed sex bias under β*2*Cas9 expression is attributable to prezygotic loss of X-bearing sperm, as directly demonstrated in Haber et al., 2024. The equivalence between split and combined designs validated the modular platform and allowed us to systematically address whether, and under what conditions, this prezygotic bias could be shifted to a postzygotic mechanism.

**Figure 2.**
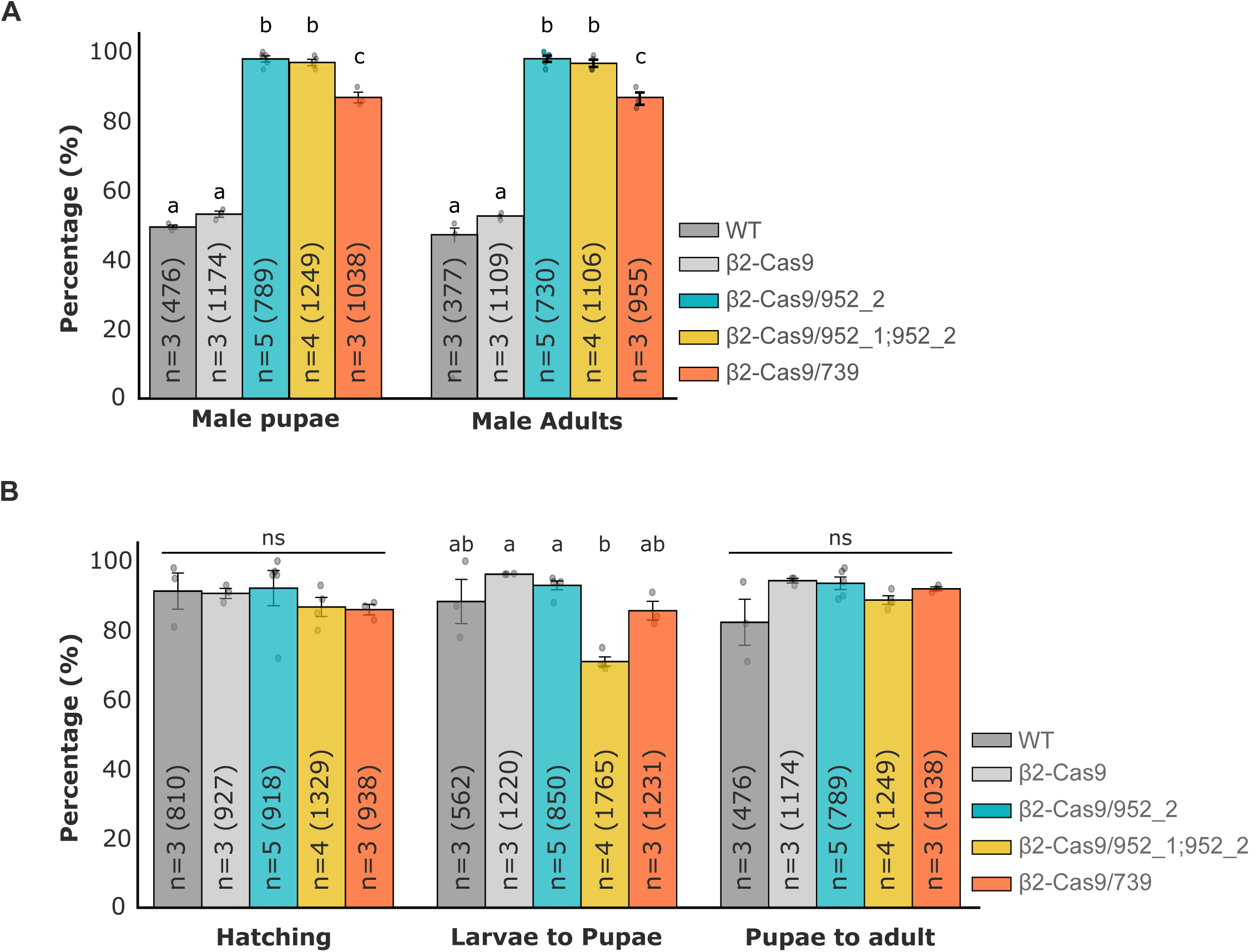
Meiotic β2Cas9 produces prezygotic sex-ratio distortion against X-linked ribosomal protein targets. (**A**) Sex-ratio distortion in F2 offspring of β2Cas9/sgRNA transheterozygous males crossed to wild-type females at pupal (left) and adult (right) stages. Five genotypes are shown: WT control, β2-Cas9 control, and experimental crosses with β2Cas9 combined with 952_2, 952_1;952_2, or 739 sgRNA lines. Numbers in bars indicate replicate number (n) and total offspring scored in parentheses. Error bars show SEM. Letters denote statistically distinct groups (Tukey’s HSD or Games-Howell post-hoc test; *p* < 0.05). (**B**) Stage-specific survival proportions (hatching, larva-to-pupa, pupa-to-adult) for the same genotypes. Comparable survival rates across all β2Cas9/sgRNA combinations and controls indicate a prezygotic rather than postzygotic mechanism of distortion. ns = not significant; letters denote statistically distinct groups where applicable.

To address this question, a second Cas9 driver line, expressing Cas9 under the control of the *zero population growth* (*zpg*) promoter, which is active in germline stem cells and spermatogonia (Tazuke *et al*., 2002; Hammond *et al*., 2021), was generated on the attP-X1 site and carrying a 3xP3:eCFP marker (**Figure 1B**). As above, *zpg*Cas9 males were then crossed to the ribosomal sgRNA strains to generate transheterozygotes for crossing to WT females. In contrast to the β*2*Cas9 crosses, *zpg*Cas9/sgRNA combinations produced pronounced and target-dependent fitness costs. *AgRpL37* targeting constructs combined with *zpg*Cas9 resulted in severe toxicity in larvae. Both *zpg*Cas9/952_2 and *zpg*Cas9/952_1;952_2 F1 transheterozygotes exhibited marked developmental asynchrony, pronounced size disparity within cohorts, and frequent *Minute*-like developmental arrest. Only 1.6% and 0.9%, respectively, survived to adulthood (**Supplementary Figure 1A**), likely due to leaky expression of *zpg*Cas9 in somatic tissues of transheterozygotes resulting in critical loss of AgRPL37.

On the other hand, the *AgRpS10* targeting construct permitted complete larval development (88% larva-to-adult viability) (**Supplementary Figure 1A**), but toxicity manifested instead at the level of reproductive function. *zpg*Cas9/739 F1 males were nearly sterile, siring only 47 viable larvae from 1,397 eggs laid by wild-type females (3% hatching rate) (**Supplementary Figure 1B**). Fertility of F1 *zpg*Cas9/739 females was also severely compromised. In two of three replicate cages (each containing 150 females), no eggs were laid. In the third cage, females produced only 45 eggs, compared to approximately 500 in the control cross. Of these, 43 hatched (96%), indicating that the defect lies primarily in fecundity rather than egg viability (**Supplementary Figure 1B**). Dissection of reproductive tissues confirmed the basis of these fertility defects. *zpg*Cas9/739 F1 males had markedly atrophied testes that lacked the synchronized spermatocysts characteristic of wild-type testes, while the accessory glands appeared normal (**Figure 3A**). In females, ovaries dissected at 24 and 48 hours post-blood feeding contained no vitellogenic follicles in either ovary at either timepoint, remaining structurally undifferentiated compared to the densely packed, yolk-loaded oocytes observed in wild type blood-fed females (**Figure 3B-C**). These phenotypes are consistent with a failure of gametogenesis arising from the loss of AgRPS10 in the germline due to early expression of Cas9 activity.

**Figure 3.**
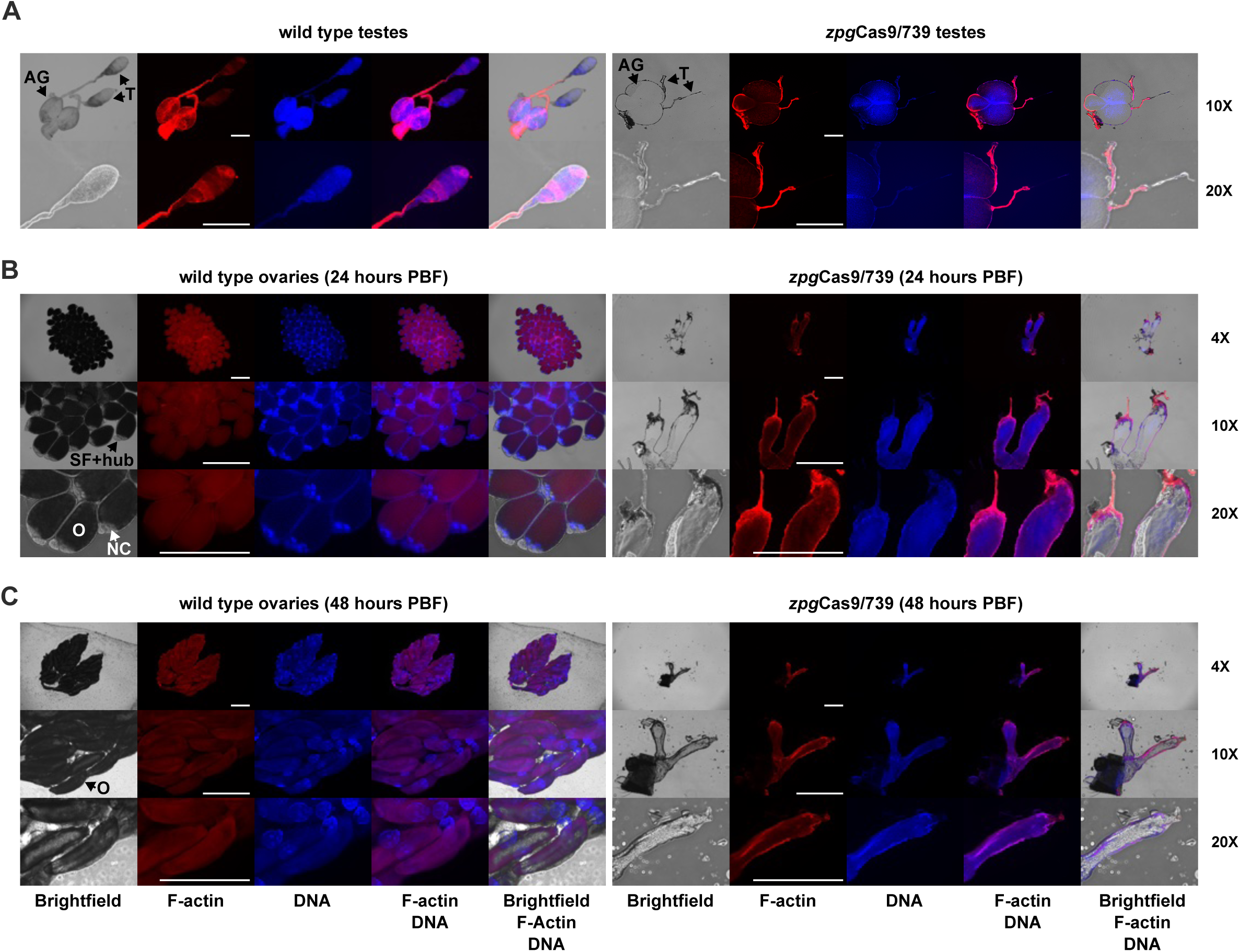
*zpg*Cas9/739 causes loss of spermatocyst development in males and failure of folliculogenesis in females. (**A**) Dissection of male reproductive tissues from wild-type (left) and *zpg*Cas9/739 F1 transheterozygous (right) males at 10X and 20X magnification. In wild-type males, accessory glands (AG) and paired testes (T) are indicated. At 20X, F-actin staining reveals synchronised spermatocysts containing cohorts of germ cells undergoing coordinated development within a shared cyst envelope. In *zpg*Cas9/739 males, accessory glands (AG) remain intact while the testes (T) are markedly atrophied and devoid of spermatocysts. (**B-C**) Ovaries dissected from wild-type (left) and *zpg*Cas9/739 F1 females (right) at 24 hours (**B**) and 48 hours (**C**) post-blood feeding (PBF), imaged at 4X, 10X, and 20X. Wild-type ovaries at 24 hours PBF contain normally developing follicles containing maturing oocytes (O). Also indicated are secondary follicles and attached hubs (SF+hub) at the apical tip of each ovariole, and nurse cells (NC) within the primary follicle containing the oocyte. By 48 hours PBF, wild-type ovaries are packed with large oocytes (O). In *zpg*Cas9/739 females, ovarioles remain structurally undifferentiated at both timepoints, with no detectable follicle development at either 24 or 48 hours PBF. Channels: brightfield, F-actin (phalloidin stain, red), DNA (DAPI stain, blue), and merged overlays as indicated. Scale bars, 500 μm (shown in F-actin panels).

### The X-linked gene *wupA* supports sex-ratio distortion under distinct germline Cas9 drivers

Given that *zpg*-driven Cas9 expression was incompatible with ubiquitously required ribosomal protein genes, we next asked whether a more functionally restricted X-linked target could support sex-ratio distortion without imposing toxicity on transheterozygous males. We therefore selected AGAP001053, the *An. gambiae* ortholog of *wings up A* (*wupA,* previously *heldup*), which encodes Troponin I, a conserved sarcomeric thin-filament protein required for muscle contraction and the only non-ribosomal protein haplolethal gene on the X-chromosome of *D. melanogaster* (Ferrus *et al*., 1990; Barbas *et al*., 1991; Beall and Fyrberg, 1991; Prado, Canal and Ferrús, 1999; Casas-Tintó and Ferrús, 2021). In *D. melanogaster*, pre-meiotic targeting of *wupA* using *nanos-*driven Cas9 produced strong male-biased sex ratios through postzygotic female lethality (Lawler *et al*., 2024), making its mosquito ortholog an attractive candidate. The *wupA* locus is located on the X chromosome between *RpL37* and the 28S rDNA cluster at position 21.3 Mbp (**Figure 1A**) and encodes eight putative alternatively spliced isoforms (**Supplementary Figure 2A**). According to RNAseq data (Papa *et al*., 2017), *wupA* expression is first detectable in 24-hour-old eggs, increasing strongly through larval development and peaking during the L2-3 larval instar, before declining progressively through late larval and adult stages (**Supplementary Figure 2B**). To disrupt all isoforms simultaneously, we generated a construct containing two sgRNAs targeting exons 2 and 5, which are shared across all predicted transcripts (**Supplementary Figure 2A**), and integrated it into the attP-E docking site on chromosome 3 (**Figure 1B**).

We first evaluated whether targeting *wupA* produces sex-ratio distortion when combined with either the *zpg* or β*2-tubulin* Cas9 driver. Among F2 progeny of β*2*Cas9/*wupA* males crossed to wild-type females, 98% of pupae and 99% of adults were male. *zpg*Cas9/*wupA* crosses also produced significant male bias, with 68% male pupae (one-way ANOVA: *p <* 0.0001) and 89% male adults (Welch’s one-way ANOVA: *p <* 0.0001), whereas all control crosses yielded approximately equal sex ratios (46-53%) (**Figure 4A**). To distinguish pre- from postzygotic effects, we quantified stage-specific survival across development. Egg hatching rates did not differ significantly between experimental and control groups (one-way ANOVA: *p = 0.2684*). In the β*2*Cas9/*wupA* cross, larval-to-pupal and pupal-to-adult survival were comparable to the β*2*Cas9-only and WT control, consistent with a predominantly prezygotic mechanism of distortion similar to that observed with ribosomal protein targets (**Figure 4B**). By contrast, F2 offspring from the *zpg*Cas9/wupA cross showed significantly reduced larval-to-pupal survival (64%, vs 92% in the *zpg*Cas9 control; one-way ANOVA: p = 0.0007) and pupal-to-adult survival (69%, vs 98% in the *zpg*Cas9 control; Welch’s one-way ANOVA: p = 0.0187; **Figure 4B**), consistent with postzygotic female-specific lethality.

**Figure 4.**
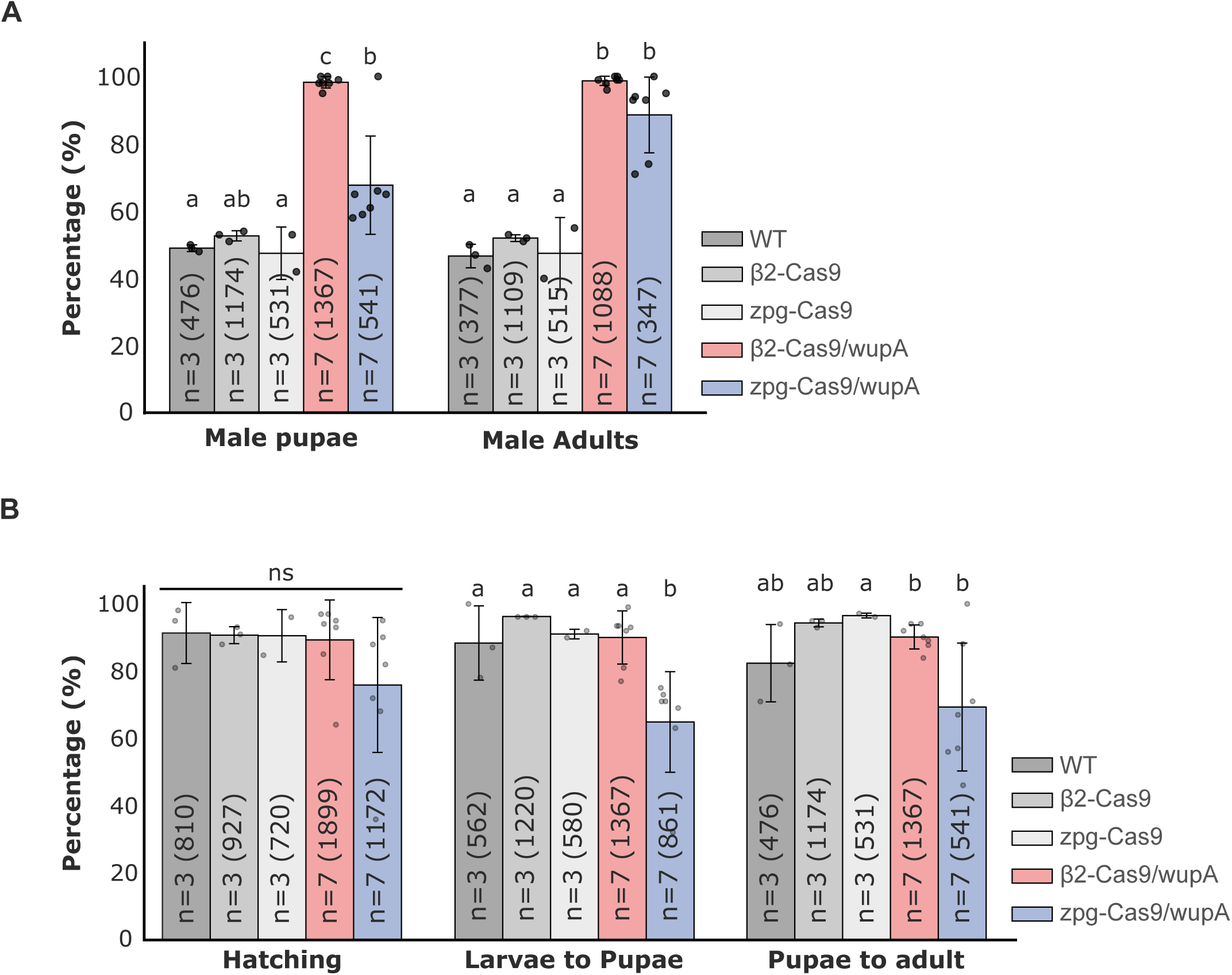
*wupA* targeting produces driver-dependent sex-ratio distortion through distinct mechanisms. (**A**) Sex ratio (% male) at pupal and adult stages in crosses between *wupA* sgRNA females and β2Cas9 or zpgCas9 males. (**B**) Developmental metrics (hatching rate, larval-to-pupal survival, and pupal-to-adult survival) across genotypes. Reduced larval-to-pupal and pupal-to-adult survival is evident only in the *zpg*Cas9/*wupA* cross, consistent with postzygotic female lethality. Different letters indicate significant pairwise differences (p < 0.05, Tukey HSD or Games-Howell post-hoc, as appropriate)

Close examination of developing F2 progeny at the pupal-adult transition revealed a striking spectrum of female-specific phenotypes in *zpg*Cas9/*wupA* crosses, consistent with progressive failure of muscle functions required for eclosion and flight (**Figure 5A-B**). Of female pupae scored, 41% drowned without beginning eclosion, while a further 33.8% initiated but failed to complete emergence, 26.6% arresting at early stages and 7.2% at late stages of eclosion. Among those that did fully escape the pupal casing, 25.2% were flightless adults and unable to leave the water surface of the pupal bowl. Altogether, only 35 of 861 larvae (4%) produced daughters capable of flight, accounting for the pronounced male bias observed among adults (flightless females were not counted as viable female adults). This phenotypic profile is consistent with the essential role of *wupA* in muscle function, and suggests that disruption of the paternal *wupA* copy in females is sufficient to impair the musculature required for eclosion and flight. To investigate potential morphological defects underlying this phenotype, we performed X-ray micro-computed tomographic (microCT) imaging on flightless females from *zpg*Cas9/*wupA* crosses and compared them to wild-type controls (**Figure 5C**). The three principal flight muscle groups, one set of dorsal longitudinal muscles (DLMs) and two sets of dorsoventral muscles (DVMs), were morphologically indistinguishable between flightless females and wild-type controls at the macroscopic level. However, scan resolution was insufficient to resolve fine structural details of the axillary sclerites and their associated musculature. Defects in sclerite-muscle coupling could therefore account for the flightless and eclosion-defective phenotypes without producing detectable changes in bulk flight muscle morphology.

**Figure 5.**
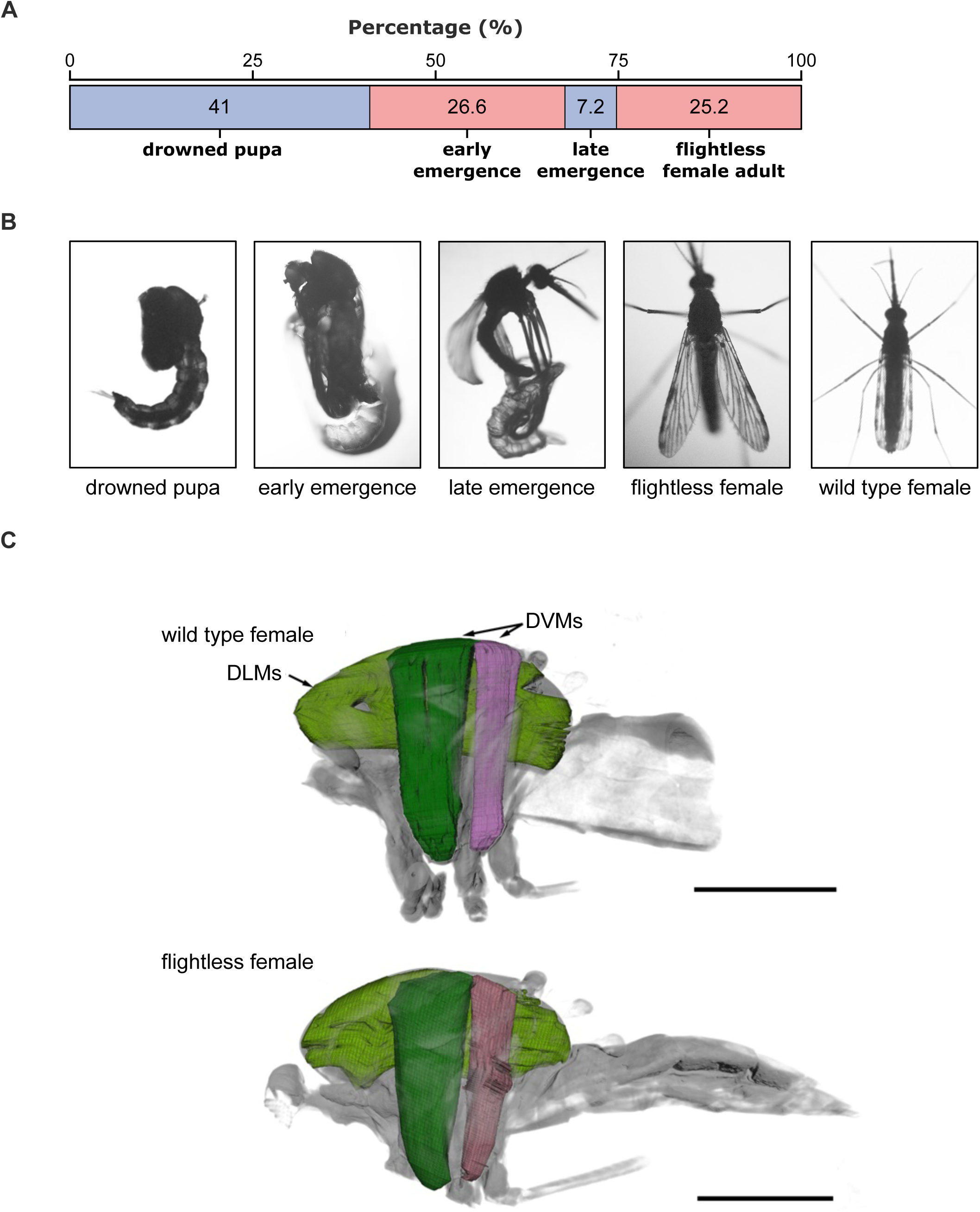
*wupA* targeting impairs eclosion and flight capacity in SRD daughters. (**A**) Breakdown of female eclosion outcomes in the *zpg*Cas9/wupA cross, expressed as percentages of 139 female pupae scored. Categories represent developmental arrest stages (drowned pupae, early emergence arrest, late emergence arrest) and flightless adult. (**B**) Representative images showing each phenotypic outcome with a wild type female included for comparison. (**C**) Comparison of flight muscle morphology between wild type females and flightless daughters of *zpg*Cas9/*wupA* males using microCT imaging. Principal flight muscles are visualized: dorsal longitudinal muscles (DLMs, light green) and two sets of dorsoventral muscles (DVMs - dark green and pink). Muscle morphology was similar between flight-capable and flightless females, indicating that *wupA*-induced flightlessness does not result from macroscopic defects in muscle architecture. The scale bar represents 650 μm.

In summary, these findings demonstrate that *wupA* targeting yields Cas9 driver-dependent distortion via distinct outcomes: under meiotic β*2*-driven expression, sex bias is prezygotic, whereas under early, pre-meiotic *zpg*-driven expression, distortion arises through female-specific post-embryonic lethality following inheritance of disrupted X-linked alleles, as originally designed for X-poisoning.

### Female-specific, post-embryonic lethality underlies *zpg*Cas9/*wupA* distortion

The preceding experiments indicated that *zpg*Cas9-mediated *wupA* disruption results in sex distortion via offspring lethality during development, but did not directly demonstrate that sex chromosome transmission is unaffected or that lethality is specific to daughters. To resolve this, we tracked offspring sex ratios across development using a Y-linked 3xP3-RFP fluorescent marker (called Y^RFP^ below) (Bernardini *et al*., 2014). Y^RFP^ homozygous *zpg*Cas9 males were crossed to *wupA*-sgRNA females to generate transheterozygous males (Y^RFP^/*zpg*Cas9/*wupA*), which were then backcrossed to wild-type females (**Figure 6A**). As a control, Y^RFP^/*zpg*Cas9 males were crossed to wild-type females.

**Figure 6.**
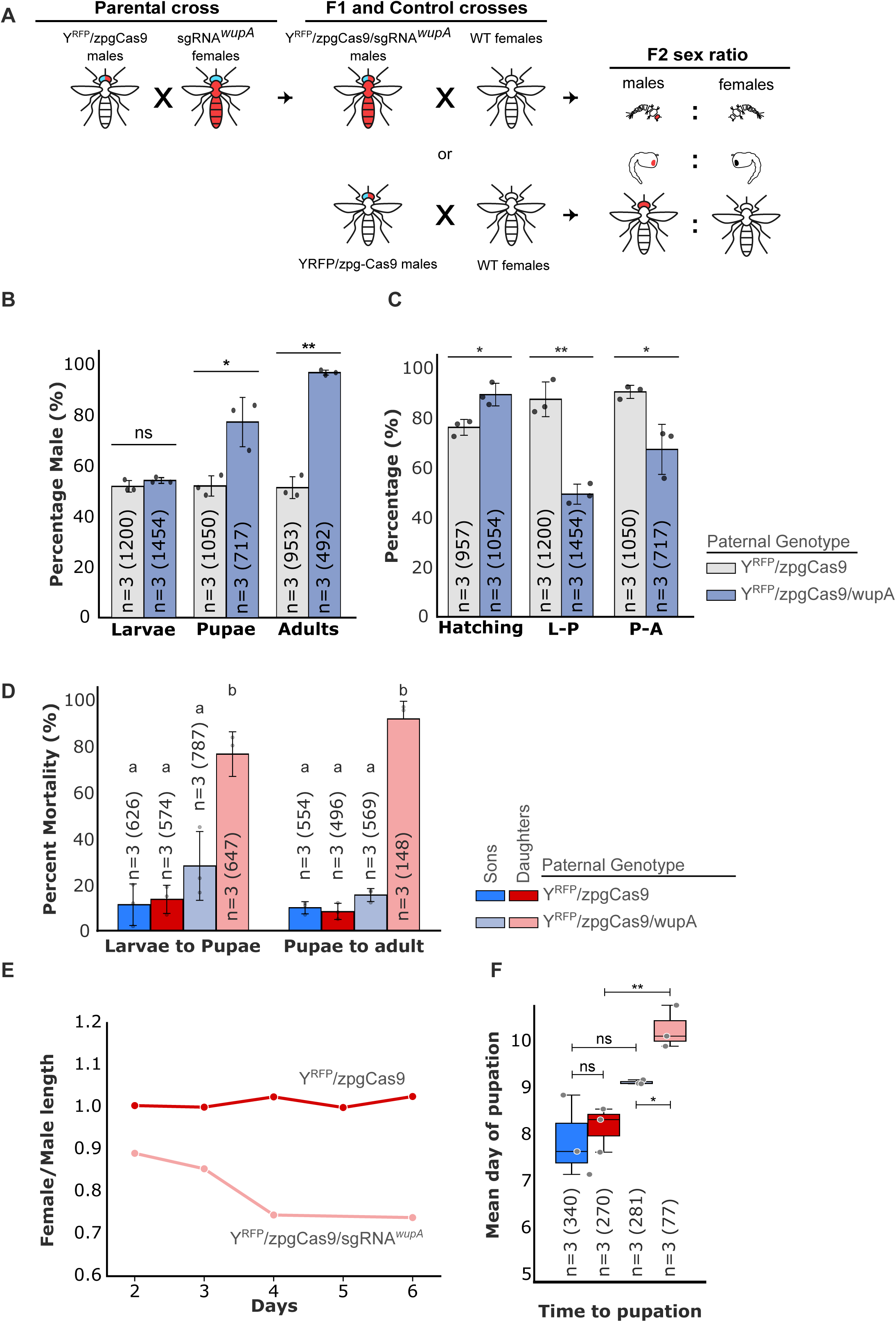
Y-linked fluorescent marker tracking confirms female-specific postzygotic lethality in zpgCas9/wupA crosses. (**A**) Crossing scheme. Homozygous Y^RFP^/*zpg*Cas9 males (carrying both a Y-linked 3xP3:RFP fluorescent marker and the *zpg*-Cas9 transgene) were crossed to *wupA* sgRNA females to generate Y^RFP^/*zpg*Cas9/*wupA* transheterozygous males, which were backcrossed to wild-type females. Y^RFP^/*zpg*Cas9 males crossed to wild-type females served as controls. Offspring sex was screened at the L1 stage based on the 3xP3:RFP fluorescence. (**B**) Percentage of male offspring at larval, pupal, and adult stages for experimental and control crosses. Male bias increases progressively across developmental stages in the experimental cross only. Error bars, SEM; asterisks indicate significant pairwise differences based on two-tailed Welch’s *t*-tests (*, *p* < 0.05; **, *p* < 0.01). (**C**) Hatching rate and overall survival (larva-to-pupa, L-P; pupa-to-adult, P-A) for both genotypes. Hatching rate was significantly lower in the control cross, but in later stages L-P and P-A survival was significantly lower in the experimental cross. (**D**) Sex-specific percent mortality at larval-to-pupal and pupal-to-adult transitions by sex and paternal genotype. Daughter mortality in the zpgCas9/wupA cross (light pink bars) is significantly elevated relative to all other groups at both developmental stages. (**E**) Female-to-male larval length ratio across developmental days 2-6 for Y^RFP^/*zpg*Cas9/*wupA* (pink line) and Y^RFP^/*zpg*Cas9 control (red line) crosses. Daughters in the experimental cross are consistently smaller than their male siblings from day 2 onward, with the ratio declining to ∼0.74 by days 4-6. (**F**) Mean day of pupation for male and female offspring of control and experimental crosses. Female pupation is significantly delayed in the Y^RFP^/*zpg*Cas9/*wupA* cross relative to controls and to male siblings.

The sex ratio among newly hatched L1 larvae did not differ between Y^RFP^/*zpg*Cas9/*wupA* and the Y^RFP^/*zpg*Cas9-only control, confirming that X-chromosome transmission is not compromised by *zpg*Cas9-mediated *wupA* disruption (**Figure 6B**). Male bias increased significantly in subsequent stages, reaching 78% at the pupal stage and 98% among emerging adults (Welch’s *t*-test, *p* = 0.0318 and *p* = 0.002, respectively; **Figure 6B**), which was associated with reduced larval-to-pupal and pupal-to-adult survival (Welch’s t-tests *p =* 0.0030 and *p =* 0.0494, respectively; **Figure 6C**). This lethality was overwhelmingly female-specific: 77% of daughters died during larval stages, and mortality of daughters reached 92% during the pupal-to-adult transition (one-way ANOVA; larva-to-pupa: *p =* 0.0002; pupa-to-adult: *p <* 0.0001; **Figure 6D**). Mortality among sons and among both sexes in control crosses remained low and did not significantly differ (**Figure 6D**). Moreover, tracking of relative larval size throughout development revealed that daughters of *zpg*Cas9/*wupA* fathers were consistently smaller than their male siblings, whereas no size differences were observed between sons and daughters of the control cross (**Figure 6E, Supplementary Figure 3**). This was reflected in a significant delay in average time to pupation: daughters of Y^RFP^/*zpg*Cas9/*wupA* males pupated on average ∼2 days later than control daughters and ∼1 day later than their experimental male siblings (Welch’s t-test; *p* = 0.0055 and *p* = 0.048, respectively; **Figure 6F**). Conversely, sons of Y^RFP^/*zpg*Cas9/*wupA* males did not differ from control (**Figure 6F**).

## Discussion

Our results show that the outcome of CRISPR targeting of X-linked genes in *Anopheles gambiae* is determined primarily by the spermatogenic stage at which Cas9 is expressed rather than by target gene identity or target-site copy number. Using a systematic comparison of Cas9 drivers and target genes in a standardized split-system design, we found that meiotic β*2-tubulin*-driven Cas9 consistently produced prezygotic sex-ratio distortion, consistent with reduced fertilization success of X-bearing sperm (Galizi *et al*., 2014; Haghighat-Khah *et al*., 2020; Haber *et al*., 2024). In contrast, shifting Cas9 activity to earlier spermatogenic stages produced strongly target-dependent outcomes: targeting ribosomal protein genes caused severe developmental or reproductive toxicity in transheterozygous carriers, whereas targeting *wupA*, which appears not to be required in the male germline for fertility, enabled the intended postzygotic lethality of daughters, characteristic of X-poisoning.

These findings resolve a central question raised from our previous work, in which attempts to engineer X-poisoning in *An. gambiae* consistently yielded prezygotic distortion rather than postzygotic lethality of daughters (Haber *et al*., 2024). At the time, the expectation was that targeting single-copy haploinsufficient genes rather than repetitive X-shredding-like sites would bypass segregation defects and allow transmission of mutant alleles to daughters, as we first observed in the fly model (Fasulo *et al*., 2020). Our results now show that this expectation does not hold in mosquitoes under meiotic Cas9 activity, where X-chromosome cleavage instead leads to prezygotic distortion. By contrast, earlier Cas9 activity allows transmission of disrupted X-linked alleles and enables postzygotic daughter lethality.

The consistent prezygotic outcome across all β*2*Cas9 combinations indicates that meiotic Cas9 activity imposes a fixed outcome on X-linked targeting, irrespective of target gene identity, sgRNA design, or target-site number. This held across both ribosomal protein genes (*AgRpS10*, *AgRpL37*) and the muscle gene *wupA*, regardless of the number of guide RNA target sites. Notably, *wupA* targeting under β*2*Cas9 produced 99% males without any detectable reduction in offspring survival, demonstrating that targets capable of producing postzygotic lethality instead yield prezygotic distortion when Cas9 is active during meiosis.

A plausible explanation for the timing-dependent influence of Cas9 activity on SRD outcomes is that DNA breaks induced during meiosis trigger elimination of damaged X-bearing sperm or impair their ability to successfully fertilize. In contrast, cleavage occurring earlier in germline development may allow repair through mutagenic end-joining while preserving sperm viability, enabling disrupted paternal X chromosomes to be transmitted to offspring. Under this model, meiotic cleavage effectively converts diverse X-chromosome targets into functional X-shredders by selectively favoring the transmission of untargeted Y-bearing sperm, whereas earlier cleavage allows mutant alleles to segregate normally and manifest as postzygotic phenotypes in daughters. Although the precise cellular mechanisms remain to be determined, these results indicate that the spermatogenic stage of Cas9 activity is a key determinant of whether X-linked cleavage results in segregation distortion or postzygotic lethality.

The fitness costs observed when *zpg*-driven Cas9 was combined with ribosomal protein targets reflect the essential roles of these genes in both germline and somatic cells. The distinct phenotypic consequences observed for each target likely reflect differences in target gene expression levels and/or functional requirements across tissues. *AgRpL37* targeting caused severe, *Minute*-like larval developmental defects, suggesting that low levels of Cas9 expression outside the germline are sufficient to induce organismal toxicity. This somatic leakiness of the zpg promoter has also been reported in previous studies (Hammond *et al*., 2021). Haploinsufficiency of large-subunit ribosomal proteins is well known to induce the *Minute* phenotype in *D. melanogaster* through cellular stress responses to ribosomal imbalance (Marygold *et al*., 2007). Our previous RNA-seq analysis of X-linked ribosomal protein genes showed that *AgRpL37* is expressed at lower levels than *AgRpS10* (Haber *et al*., 2024), which may render it more sensitive to partial loss of function following disruption of a single allele. By contrast, leaky targeting of *AgRpS10* did not impair somatic development but abolished fertility in both sexes, indicating an essential role in gametogenesis. It is possible that the higher expression of *AgRpS10* in somatic tissues buffers against partial loss of function. However, its strict requirement in the germline makes it highly sensitive to disruption by *zpg*-driven Cas9 activity in germline stem cells and spermatogonia. *AgRpL37* is also likely required in the germline, but lethality of transheterozygotes prevented direct assessment of this possibility. In both cases, disruption of these ribosomal protein genes proved incompatible with *zpg-*driven Cas9 expression, indicating that broadly expressed haploinsufficient genes are unsuitable targets for X-poisoning in *An. gambiae* under early germline Cas9 activity, as disruption in either somatic or germline contexts leads to severe fitness costs. We observed a similar sterility phenotype in *D. melanogaster* when *RpS6*-targeting sgRNAs were combined with early germline *nanos*-Cas9 expression, resulting in complete male sterility (Fasulo *et al*., 2020), consistent with disruption of ribosomal protein function during spermatogenesis.

Disruption of *wupA* under pre-meiotic Cas9 expression produced a strong male bias. Y-linked fluorescent marker tracking demonstrated that this bias arises from postzygotic lethality of daughters. Sex ratios among newly hatched L1 larvae were balanced and hatching rates were higher for the *zpg*Cas9/*wupA* cross than the control, demonstrating that sex chromosome segregation is intact and that daughter mortality accumulates after the egg stage, from the first larval instar through pupation. The lethality observed in daughters confirms the haplolethal character of *wupA*: daughters inheriting a disrupted paternal X chromosome alongside the maternally derived wild-type allele lack sufficient functional Troponin I.

Unlike *D. melanogaster*, where *wupA* targeting results in female lethality predominantly during the egg stage (Lawler *et al*., 2024), in mosquitoes daughters complete embryogenesis normally but die during larval and pupal development or emerge as flightless adults. This likely reflects species-specific differences in *wupA* function across development. Importantly, this shift in the timing of lethality may influence population suppression outcomes through interactions with density-dependent larval competition, meaning it could be advantageous for mosquito genetic control. Mosquito larvae develop in discrete, resource-limited breeding sites where density-dependent mortality is a major regulator of population size. Both experimental and modelling studies have shown that the timing of mortality relative to the density-dependent phase is critical: early-acting female mortality can relax competition and partially compensate population losses, whereas lethality occurring during larval stages can interact with density-dependent processes and enhance suppression (Phuc *et al*., 2007; Yakob, Alphey and Bonsall, 2008; Vella, Gould and Lloyd, 2021; Butler and Lloyd, 2025). The progressive larval and pupal lethality we observed here therefore positions *wupA*-based X-poisoning to exploit these competition dynamics, potentially enhancing its impact as a population suppression tool in the field.

These findings have direct implications for vector control. Self-limiting Y-linked sex-ratio distortion systems differ fundamentally from invasive meiotic drives: their frequency in the population remains proportional to the release ratio, allowing the extent and duration of any population intervention to be calibrated to a release program (Burt and Deredec, 2018). This property may facilitate regulatory approval and community acceptance, particularly for early-phase field trials where geographic containment and reversibility are priorities. Beyond standalone deployment, such systems offer versatility as combination tools, for example to prime populations prior to introduction of more persistent, self-sustaining gene drive, or to enable spatially targeted interventions across regions with differing regulatory frameworks. The demonstration that *wupA*-based female-specific lethality is achievable in *An. gambiae* therefore provides a concrete foundation for developing Y-linked self-limiting SRD systems for malaria vector control.

A key challenge for implementing a Y-linked X-poisoning system is transcriptional silencing of Y chromosome transgenes. Encouragingly, multiple lines of evidence indicate that pre-meiotic germline promoters can escape this silencing. Y-linked transgenes driven by the *vasa* promoter retain functional expression in *D. melanogaster* (Gamez *et al*., 2021) and in *An. gambiae* (Bernardini *et al*., 2014; Tolosana *et al*., 2025), consistent with the idea that promoters active prior to meiotic sex chromosome inactivation (MSCI) may have access to a more permissive chromatin environment. The pre-meiotic *zpg* promoter used here belongs to this class, making it a candidate for driving Y-linked Cas9 expression. In contrast, meiotic promoters are consistently silenced on the Y: β*2*-tubulin-driven transgenes produce little to no detectable expression in *An. gambiae* testes (Alcalay *et al*., 2021) and on the Y of *D. melanogaster* (Arien *et al*., 2025). However, pre-meiotic activity alone does not guarantee Y-linked expression. In *Drosophila suzukii*, Y-linked *nanos*-Cas9 expression is substantially reduced relative to autosomal controls, with substantial Cas9 activity observed only in one case of anomalous double integration (Hewawasam, Yamamoto and Scott, 2025). Together, these findings indicate that expression of transgenes on the Y chromosome may be promoter-dependent and cannot be predicted solely by pre- versus post-meiotic activity. Identifying the promoter and integration contexts that support robust Y-linked expression therefore remains a critical step toward functional self-limiting sex-ratio distortion systems. The identification of *wupA* as an effective X-poisoning target in both flies and mosquitoes suggests that tissue-restricted X-linked haploinsufficient genes may represent a broadly applicable class of targets for postzygotic sex-ratio distortion across dipteran species, expanding the genetic toolkit available for vector and pest control.

## Methods

### Mosquito rearing

*Anopheles gambiae* mosquitoes were maintained at 28 °C, 80% relative humidity, under a 12 h light/12 h dark cycle. Strains used included the G3 wild-type laboratory strain (Imperial College London), the Y-linked 3xP3-RFP strain attP-Y (Bernardini *et al*., 2014), and two autosomal attP docking lines – attP-X1 and attP-E (IBMC, Strasbourg University; Meredith *et al*., 2011; Volohonsky *et al*., 2015). Larvae were reared in demineralized water at densities of 200-250 individuals per 500 mL tray and fed daily with ground commercial fish food. Adults were maintained on 10% sucrose supplemented with 1% nipagin *ad libitum*. Adults were allowed to mate for 4-7 days and were blood-fed on bovine blood using either a Hemotek membrane feeder (Hemotek Ltd.) or the 3D-Feedy system (Diptera.ai) (Ens *et al*., 2025). Oviposition bowls were provided 72 h after blood feeding, composed of a Whatman 3 filter paper embedded in demineralized water. Newly hatched L1 larvae were transferred to fresh trays at 48 h and reared until pupation.

### Cloning of transformation plasmids

Constructs were assembled by Gibson assembly (NEBuilder HiFi DNA Assembly, NEB) or Golden Gate assembly. PCR reactions used Q5 Hot Start High-Fidelity 2× Master Mix (NEB). All plasmids were verified by Sanger sequencing (**Supplementary Materials**). To establish the split CRISPR-Cas9 system we generated two sets of plasmids for either Cas9 or gRNA components, using distinct fluorescent markers for each construct. Construction of both sets was based on the same plasmid backbone containing a single phiC31 attB site to facilitate site-specific integration and a fluorescent marker. The β*2*-Cas9 and *zpg*-Cas9 constructs were marked with *3xP3*-RFP and 3XP3:CFP, respectively. For sgRNA constructs, the Pol III promoter of *U6* (AGAP013695) was used to drive expression of each sgRNA. The 739 sgRNA construct was labelled with an *D. melanogaster Actin5C-*eGFP marker, while *Actin5C*-RFP was used for the 952_2, 952_1;952_2, and *wupA* sgRNAs constructs. The *wupA* (AGAP001053) target sites were selected in exons 2 and 5, which are present in all isoforms using the Doench on-target activity score (0.537 and 0.743, respectively) and the Zhang specificity score (100% specificity with no off-targets at 0-2 mismatches; 91.08% specificity allowing up to three mismatches, with 40 predicted off-target sites). No off-target sites were located within coding sequences. sgRNA targeting ribosomal protein genes *AgRpS10* (AGAP000739) and *AgRpL37* (AGAP000952) were taken from Haber et al. (2024). sgRNA sequences used for all targets are provided in **Supplementary Table 1**.

### Embryo microinjection and transgenesis

Embryo microinjections were performed as described in (Volohonsky *et al*., 2015) using an Eppendorf FemtoJet 4x injector and a Narishige MM-94 micromanipulator mounted on an Olympus IX53 inverted microscope. To generate transgenic *Anopheles gambiae* strains expressing sgRNAs, 1-2 hours old embryos from the attP-E docking line were injected with plasmid mixtures containing transformation constructs and *vasa*:phiC31 integrase helper plasmid at a 2:1 ratio and total DNA concentrations of 400-800 μg/µl. Injected G0 larvae were reared to adulthood and outcrossed to wild-type mosquitoes. G1 progeny were screened as larvae and sorted by fluorescent marker: Actin5C-GFP for the 739 sgRNA line and Actin5C-RFP for the 952_1, 952_2, 952_1;952_2, and *wupA* sgRNA lines. Positives were reared to adulthood and intercrossed to establish homozygous stocks. *zpg*-Cas9 and β*2*-Cas9 lines were generated by co-injection of a mix of *zpg*-Cas9 and β*2*-Cas9 plasmids together with a *vasa*:phiC31 integrase helper plasmid, at final concentrations of 330 μg/μl for each of the transformation plasmids and 80 μg/μl for the helper plasmid. G0 individuals exhibited transient expression of all fluorescent markers – 3xP3:CFP for *zpg*-Cas9 and 3xP3:RFP for β*2*-Cas9. Pupae were sexed and outcrossed to attP-X1 docking-line individuals of the opposite sex. Fluorescent G1 progeny were screened, intercrossed to homozygosity over multiple generations. Homozygous populations were established and sorted using a Complex Object Parametric Analyzer and Sorter (COPAS - Union Biometrica).

### Sex-ratio distortion and developmental assays

To generate individuals expressing both Cas9 and sgRNA components, *zpg*Cas9 or β*2*Cas9 males were crossed to females carrying 739, 952_1;952_2, 952_2, or *wupA* sgRNA constructs. From each Cas9/sgRNA cross, 30–40 transheterozygous F1 males were mated to 40–50 wild-type females. Oviposition cups were provided 48 hrs post-blood-feeding and left overnight. L1 larvae were transferred 48 hrs after egg laying to fresh trays at a density of 200–250 larvae per tray in 500 mL of demineralized water. Developmental progression from egg to adult was monitored daily.

Hatching rate was determined by scoring the number of hatched larvae relative to the total number of eggs laid. Pupation and adult eclosion were recorded daily by direct counting of pupae and newly emerged adults, respectively. To determine L1 sex ratios in *zpg*Cas9 crosses carrying the male-specific Y^RFP^ marker, 400–500 larvae were screened 48 hrs after egg laying for 3xP3-RFP fluorescence, and sex ratio was scored.

### Tracking of larval size in offspring of Y^RFP^/*zpg*Cas9/*wupA* males

40-50 Y^RFP^/*zpg*Cas9/*wupA* transheterozygous males and Y^RFP^/*zpg*Cas9-only males were separately crossed to 70-80 wild-type females. Offspring were collected and reared at a density of ∼250 larvae per tray, beginning 48 hours post-oviposition. Each day, 20 male and 20 female larvae were sexed using the Y^RFP^ fluorescent marker (3xP3-RFP). Selected larvae were transferred to a lid of a 96-well plate in a water bubble to restrict movement and maintain a two-dimensional posture. Larvae were imaged using a Thermo Fisher EVOS M5000 fluorescence microscope, and body length (head-to-tail) was measured for each individual (see example in **Supplementary Figure 3)**. Measurements were performed daily until the onset of pupation, at which point final measurements were recorded. Documented larvae were discarded. Due to excess female mortality during larval development, female offspring of Y^RFP^/*zpg*Cas9/*wupA* males became progressively less abundant over time; therefore, fewer than 20 females were measured at later time points.

### Time to pupation

40-50 Y^RFP^/*zpg*Cas9/*wupA* transheterozygous males and Y^RFP^/*zpg*Cas9-only males were crossed separately to 70–80 wild-type females. 48 hours post-oviposition L1 larvae were sexed by screening for the Y^RFP^ fluorescent marker (3xP3-RFP) and distributed into three replicate groups of 150 individuals per sex per tray. Larvae were reared under standard conditions until pupation, and the number of pupae was recorded daily for each replicate and for each sex.

### Tissue staining and imaging

Testes and ovaries were dissected in 1X Phosphate-Buffered Saline (PBS) and fixed for 30 minutes at room temperature in 4% paraformaldehyde (Thermo fisher, 043368.9L) in 1X PBS. Tissues were permeabilized for 45 min in 1X PBS, 0.1% Triton (Sigma-Aldrich, X-100), then stained for 1 hr in 1X PBS containing Atto 594-phalloidin (Sigma-Aldrich, 51927) and l DAPI (Sigma-Aldrich, D9542) according to manufacture working solution protocols. Samples were washed twice in 1X PBS, mounted in 1X PBS, and imaged on a Thermo Fisher EVOS M5000 fluorescence microscope using appropriate filter sets for DAPI and 594-phalloidin channels.

### Sample preparation and micro-CT

Adult wild-type and flightless zpgCas9/wupA females were collected for imaging. Specimens were first decapitated and then immersed in 1% Tween-20 (Bio-Rad, 1706531) to reduce surface tension and facilitate fixative penetration. Samples were fixed for 1 hr at room temperature in 2.5% paraformaldehyde/glutaraldehyde (Electron Microscopy Sciences, 15949), followed by incubation at 4 °C for 72 h. Following fixation, specimens were washed three times in cold (4 °C) 1× PBS for 10 min each. Samples were then stained in 1% Lugol’s iodine solution (Sigma-Aldrich, L6146) at 4 °C for 48 h. Prior to scanning, specimens were rinsed in demineralized water to remove excess iodine. Micro–computed tomography (microCT) imaging was performed using a Bruker SkyScan 1272 system at the Center for Scientific Imaging (CSI) at the Faculty of Agriculture, Food and Environment of HUJI (Rehovot). Three-dimensional visualization and segmentation were performed in Amira 3D 2025 (Thermo Fisher Scientific).

### Statistical analysis

All statistical analyses were performed using JMP Pro 19. Proportional data (sex ratios, hatching rates, and survival transitions) were analysed at the replicate level using one-way ANOVA. Homogeneity of variance was assessed using Levene’s test; when violated (p < 0.05), Welch’s ANOVA with Welch–Satterthwaite correction was applied. Post-hoc pairwise comparisons were conducted using Tukey’s HSD (equal variances) or Games–Howell (unequal variances). For comparisons involving two groups only, two-tailed Welch’s t-tests were used. Statistical significance was assessed at α = 0.05. **Supplementary File 2** reports complete statistical results for all analyses.

## Author Contributions

LBL, DAH, YA and PAP conceived and designed the study. LBL, DAH, AS, ESY, EM generated transgenic constructs and strains and performed all mosquito experiments with the help of SPS and ESMK. LRD performed micro-CT analyses. PAP supervised the project, secured funding, and wrote the manuscript together with LBL and with input from all authors. All authors read and approved the final manuscript.

## Acknowledgements

We thank Doron SY Zaada for discussions, experimental suggestions and support. We are also thankful to Carmel Katz for assistance with the statistical analysis. We thank Gleb Ens, Neta Levine and Ruth Shacham for mosquito strain maintenance. We thank Einat Zelinger and Tally Kossovsky at the CSI HUJI.

## Funding

This work was supported by research grants to PAP from the Gates Foundation (INV-004363 and INV-075374) and the Israel Science Foundation (2388/19).

## Ethics

All mosquito work was conducted under institutional biosafety and arthropod containment approvals from the Hebrew University of Jerusalem. Blood feeding using bovine blood was performed in accordance with institutional animal use guidelines and under approved animal ethics protocols. All experiments involving genetically modified organisms complied with national and institutional regulations. All insect work was performed in facilities maintaining Arthropod Containment Level 2. This work received Institutional Approval and relevant authorizations from the Israel Ministry of Environmental Protection and Ministry of Agriculture (#1031/21).

## Competing interests

The authors declare no competing interests.

## Supplementary Data

**Supplementary Table 1:**
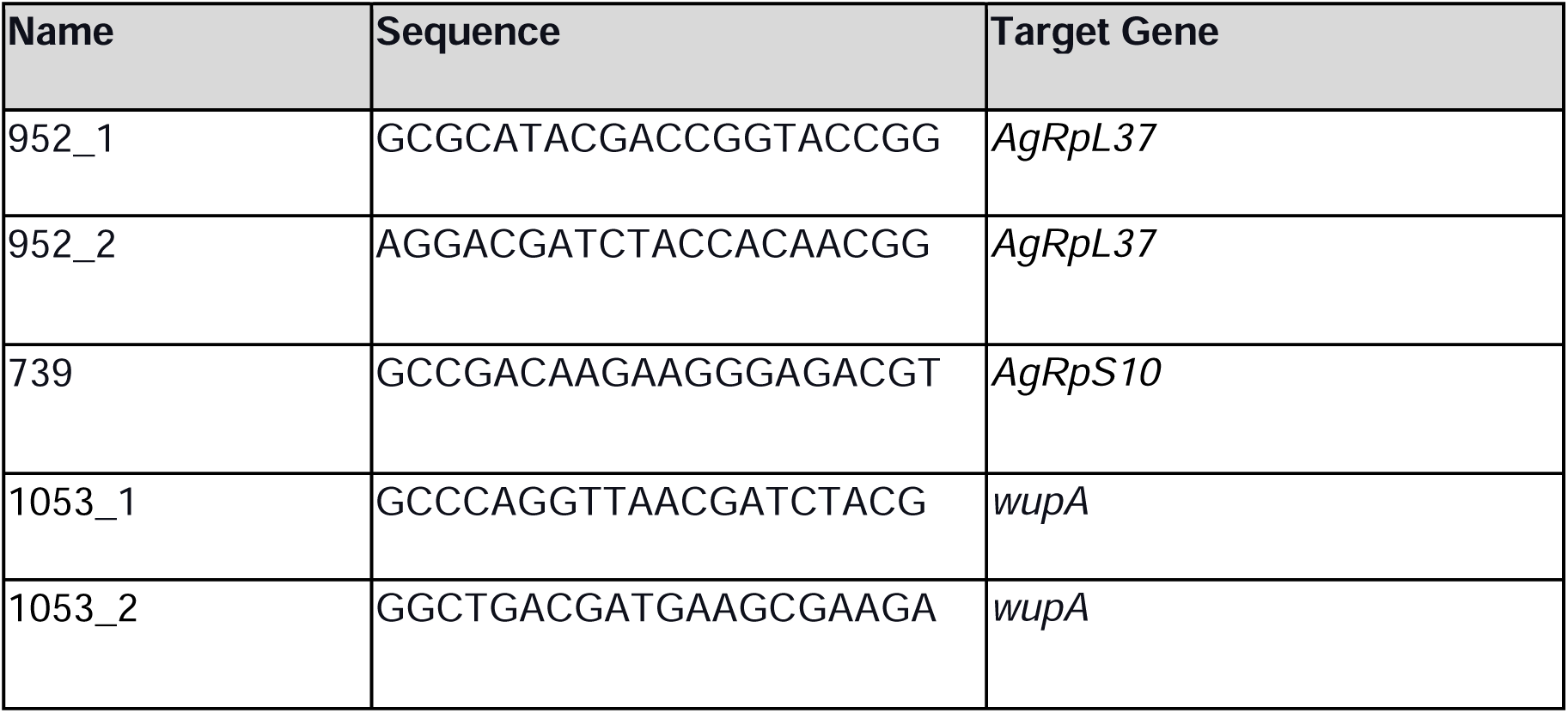
X-chromosome targeting sgRNAs.

**Supplementary File 1**: Genebank annotated plasmid sequences used in this study

**Supplementary File 2**: Statistical analyses

**Supplementary File 3**: Raw data from crosses of SRD males

**Supplementary Figure 1.**
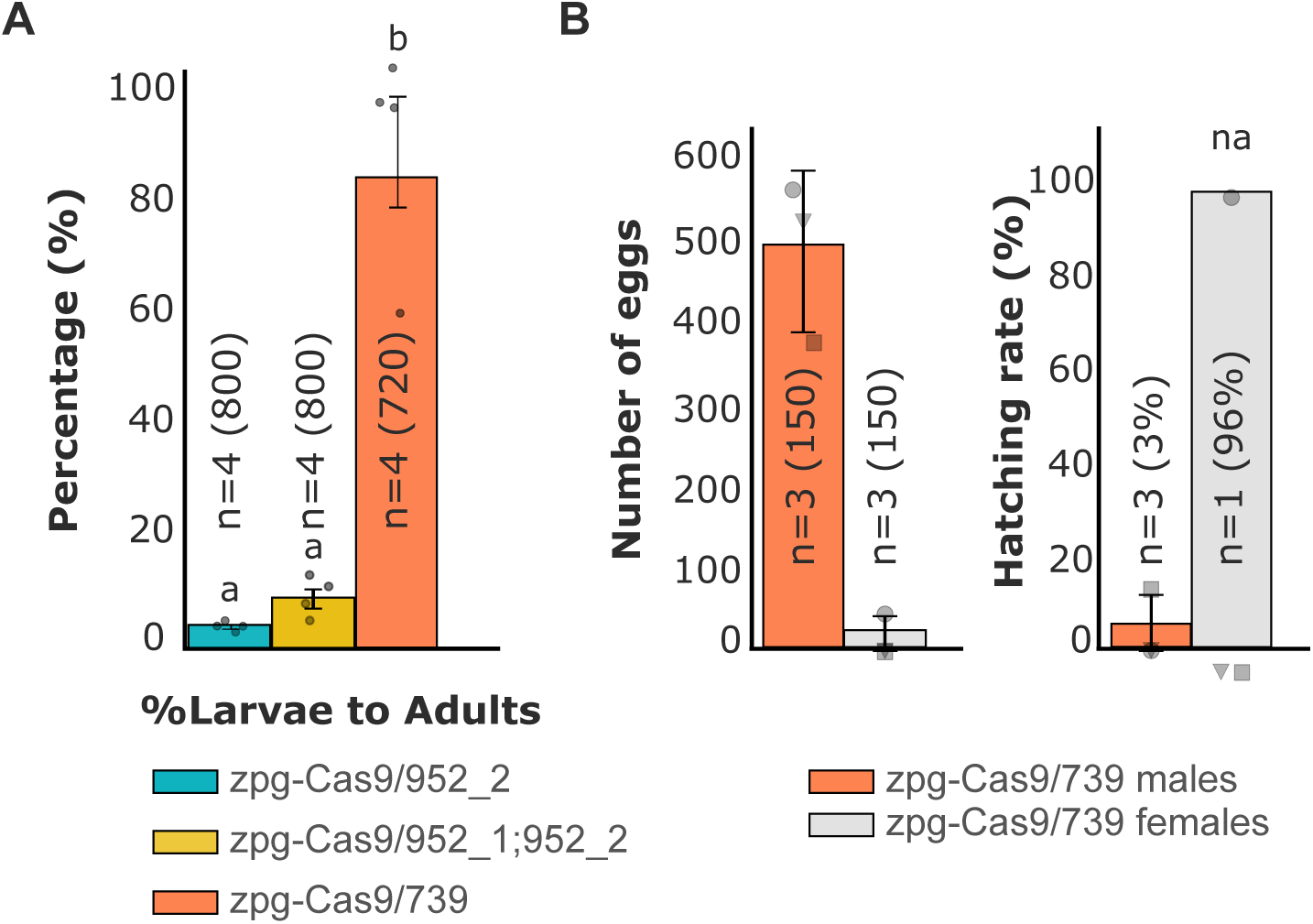
Pre-meiotic zpgCas9 combined with ribosomal protein sgRNA targets causes target-dependent fitness costs. (**A**) Percentage of larvae surviving to adulthood in F1 transheterozygotes of zpgCas9 with sgRNA targets against *AgRpL37* (952_2 and 952_1;952_2), or *AgRpS10* (739). Targeting *AgRpL37* causes near-complete larval lethality, while targeting *AgRpS10* allows high larval survival (∼85%). n=4 biological replicates; total individuals indicated in parentheses; letters denote statistically distinct groups. (**B**) Fertility of F1 zpgCas9/739 male and female transheterozygotes mated to wild-type partners. Left: number of eggs laid per replicate cage; *zpg*Cas9/739 males (orange) produce normal egg numbers when mated to wild-type females (150 females per replicate), as expected, whereas *zpg*Cas9/739 females (grey) lay very few eggs. Only one of the replicates (containing 150 females each laid eggs) laid eggs. Right: hatching rate of those eggs. Eggs sired by *zpg*Cas9/739 males show near-zero hatching (∼3%), indicating male sterility. Eggs from the one *zpg*Cas9/739 female cage had a normal hatching rate at ∼96%. Error bars, mean ± SD.

**Supplementary Figure 2.**
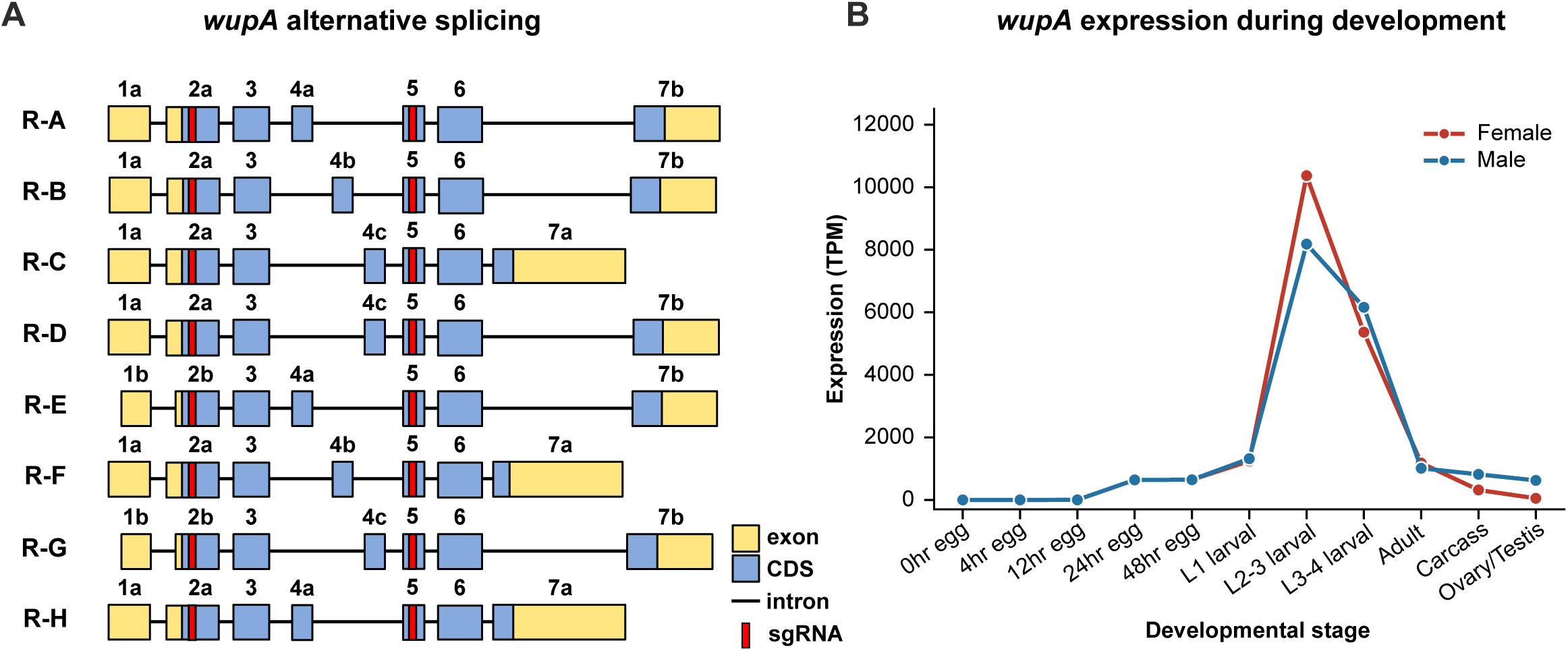
The *wupA* locus encodes eight alternatively spliced isoforms and is highly expressed during larval development. (**A**) Gene structure of *wupA* (AGAP001053) depicting eight annotated isoforms (R-A through R-H). Exons (yellow), coding sequences (blue), and introns (black lines) are shown. sgRNA target sites (red) are located within exons 2a/2b and exon 5, which are shared across all isoforms. (**B**) Expression of *wupA* across *An. gambiae* developmental stages in females (red) and males (blue), shown as transcripts per million (TPM). Expression is low in early embryos, rises through larval instars, peaks at the L2-3 larval stage in both sexes, and declines markedly in pupae and adults. Data from Papa *et al*. (Papa *et al*., 2017) and downloaded from Vectorbase.

**Supplementary Figure 3.**
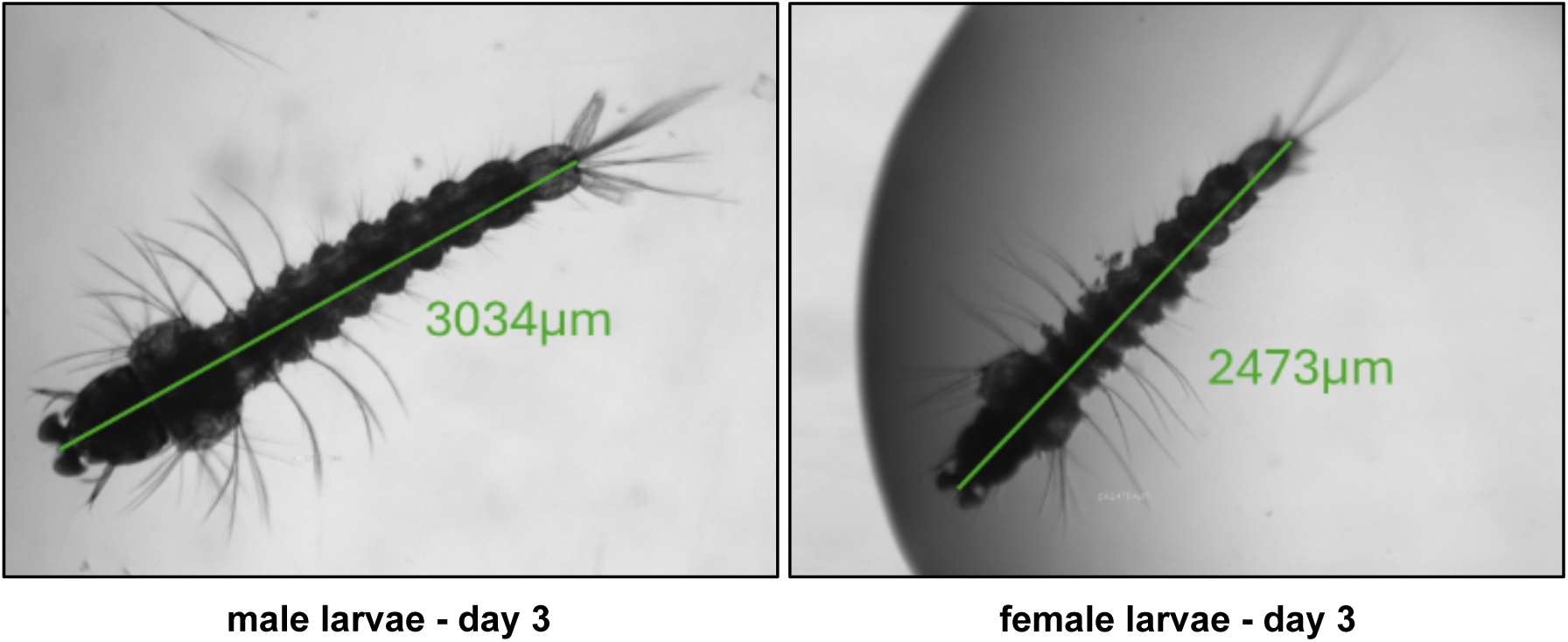
Female larvae from zpgCas9/wupA crosses are smaller than siblings. Representative larval length measurements at day 3 after hatching showing sexual dimorphism in offspring of *zpg*Cas9/*wupA* crosses.

## Notes

### Competing Interest Statement

The authors have declared no competing interest.

