## Supplementary material for "Spermatogenic context controls outcomes of engineered sex distortion in malaria mosquitoes": Supplementary File 2 - Lamdan et al 2026 - Statistics.docx

### Supplementary File 2 - Statistical Analyses

This document reports complete statistical results for all quantitative analyses in Lamdan et al. 2026, organized by figure panel. Highlighted rows in post-hoc tables indicate significant pairwise comparisons (p < 0.05). Significance codes: *** p < 0.001; ** p < 0.01; * p < 0.05; ns = not significant.

**Figure 2:**

Five genotypes: WT/WT, β2Cas9, β2Cas9/952₂, β2Cas9/952₁;952₂, β2Cas9/739. n = 3 replicates (WT/WT, β2Cas9, β2Cas9/739), n = 4 replicates (β2Cas9/952₁;952₂), n = 5 replicates (β2Cas9/952₂). All data read from CSV source. Test selection based on Levene’s test for homogeneity of variance.

### Figure 2A - % male pupae

| Group | n (replicates) | Mean | SD |
| --- | --- | --- | --- |
| WT/WT | 3 | 0.490 | 0.010 |
| β2Cas9 | 3 | 0.527 | 0.015 |
| β2Cas9/952₂ | 5 | 0.970 | 0.020 |
| β2Cas9/952₁;952₂ | 4 | 0.960 | 0.018 |
| β2Cas9/739 | 3 | 0.860 | 0.026 |

Levene's test: F(4,13) = 1.3211, p = 0.3136 ns

| Metric | Test | F | df | p |
| --- | --- | --- | --- | --- |
| % male pupae (Figure 2A) | One-way ANOVA | 536.0952 | (4, 13) | < 0.0001 *** |

Tukey HSD post-hoc pairwise comparisons:

| Group 1 | Group 2 | Mean 1 | Mean 2 | p | Sig. |
| --- | --- | --- | --- | --- | --- |
| WT/WT | β2Cas9 | 0.490 | 0.527 | 0.1846 | ns |
| WT/WT | β2Cas9/952₂ | 0.490 | 0.970 | < 0.0001 | *** |
| WT/WT | β2Cas9/952₁;952₂ | 0.490 | 0.960 | < 0.0001 | *** |
| WT/WT | β2Cas9/739 | 0.490 | 0.860 | < 0.0001 | *** |
| β2Cas9 | β2Cas9/952₂ | 0.527 | 0.970 | < 0.0001 | *** |
| β2Cas9 | β2Cas9/952₁;952₂ | 0.527 | 0.960 | < 0.0001 | *** |
| β2Cas9 | β2Cas9/739 | 0.527 | 0.860 | < 0.0001 | *** |
| β2Cas9/952₂ | β2Cas9/952₁;952₂ | 0.970 | 0.960 | 0.9300 | ns |
| β2Cas9/952₂ | β2Cas9/739 | 0.970 | 0.860 | < 0.0001 | *** |
| β2Cas9/952₁;952₂ | β2Cas9/739 | 0.960 | 0.860 | < 0.0001 | *** |

Compact letter display (groups sharing a letter are not significantly different):

| Group | Mean | Letter(s) |
| --- | --- | --- |
| WT/WT | 0.490 | a |
| β2Cas9 | 0.527 | a |
| β2Cas9/739 | 0.860 | c |
| β2Cas9/952₁;952₂ | 0.960 | b |
| β2Cas9/952₂ | 0.970 | b |

**Figure 2A - % male adults**

| Group | n (replicates) | Mean | SD |
| --- | --- | --- | --- |
| WT/WT | 3 | 0.467 | 0.035 |
| β2Cas9 | 3 | 0.520 | 0.010 |
| β2Cas9/952₂ | 5 | 0.970 | 0.020 |
| β2Cas9/952₁;952₂ | 4 | 0.957 | 0.021 |
| β2Cas9/739 | 3 | 0.857 | 0.031 |

Levene's test: F(4,13) = 1.2460, p = 0.3399 ns

| Metric | Test | F | df | p |
| --- | --- | --- | --- | --- |
| % male adults (Figure 2A) | One-way ANOVA | 359.6801 | (4, 13) | < 0.0001 *** |

Tukey HSD post-hoc pairwise comparisons:

| Group 1 | Group 2 | Mean 1 | Mean 2 | p | Sig. |
| --- | --- | --- | --- | --- | --- |
| WT/WT | β2Cas9 | 0.467 | 0.520 | 0.1018 | ns |
| WT/WT | β2Cas9/952₂ | 0.467 | 0.970 | < 0.0001 | *** |
| WT/WT | β2Cas9/952₁;952₂ | 0.467 | 0.957 | < 0.0001 | *** |
| WT/WT | β2Cas9/739 | 0.467 | 0.857 | < 0.0001 | *** |
| β2Cas9 | β2Cas9/952₂ | 0.520 | 0.970 | < 0.0001 | *** |
| β2Cas9 | β2Cas9/952₁;952₂ | 0.520 | 0.957 | < 0.0001 | *** |
| β2Cas9 | β2Cas9/739 | 0.520 | 0.857 | < 0.0001 | *** |
| β2Cas9/952₂ | β2Cas9/952₁;952₂ | 0.970 | 0.957 | 0.9318 | ns |
| β2Cas9/952₂ | β2Cas9/739 | 0.970 | 0.857 | 0.0002 | *** |
| β2Cas9/952₁;952₂ | β2Cas9/739 | 0.957 | 0.857 | 0.0008 | *** |

Compact letter display (groups sharing a letter are not significantly different):

| Group | Mean | Letter(s) |
| --- | --- | --- |
| WT/WT | 0.467 | a |
| β2Cas9 | 0.520 | a |
| β2Cas9/739 | 0.857 | c |
| β2Cas9/952₁;952₂ | 0.957 | b |
| β2Cas9/952₂ | 0.970 | b |

### Figure 2B - Hatching rates

| Group | n (replicates) | Mean | SD |
| --- | --- | --- | --- |
| WT/WT | 3 | 0.913 | 0.091 |
| β2Cas9 | 3 | 0.907 | 0.025 |
| β2Cas9/952₂ | 5 | 0.922 | 0.114 |
| β2Cas9/952₁;952₂ | 4 | 0.868 | 0.056 |
| β2Cas9/739 | 3 | 0.860 | 0.026 |

Levene's test: F(4,13) = 1.5893, p = 0.2358 ns

| Metric | Test | F | df | p |
| --- | --- | --- | --- | --- |
| Hatching rate (Figure 2B) | One-way ANOVA | 0.4747 | (4, 13) | 0.7538 ns |

Overall test not significant; no post-hoc comparisons performed.

#### Figure 2B - Larva-to-pupa survival

| Group | n (replicates) | Mean | SD |
| --- | --- | --- | --- |
| WT/WT | 3 | 0.883 | 0.111 |
| β2Cas9 | 3 | 0.962 | 0.001 |
| β2Cas9/952₂ | 5 | 0.930 | 0.028 |
| β2Cas9/952₁;952₂ | 4 | 0.710 | 0.027 |
| β2Cas9/739 | 3 | 0.857 | 0.047 |

Levene's test: F(4,13) = 3.8497, p = 0.0282 *

| Metric | Test | F | df | p |
| --- | --- | --- | --- | --- |
| Larva-to-pupa survival (Figure 2B) | Welch's one-way ANOVA | 66.2058 | (4, 5.06) | < 0.0001 *** |

Games–Howell post-hoc pairwise comparisons:

| Group 1 | Group 2 | Mean 1 | Mean 2 | p | Sig. |
| --- | --- | --- | --- | --- | --- |
| WT/WT | β2Cas9 | 0.883 | 0.962 | 0.7475 | ns |
| WT/WT | β2Cas9/952₂ | 0.883 | 0.930 | 0.9358 | ns |
| WT/WT | β2Cas9/952₁;952₂ | 0.883 | 0.710 | 0.3102 | ns |
| WT/WT | β2Cas9/739 | 0.883 | 0.857 | 0.9927 | ns |
| β2Cas9 | β2Cas9/952₂ | 0.962 | 0.930 | 0.2418 | ns |
| β2Cas9 | β2Cas9/952₁;952₂ | 0.962 | 0.710 | 0.0013 | ** |
| β2Cas9 | β2Cas9/739 | 0.962 | 0.857 | 0.1770 | ns |
| β2Cas9/952₂ | β2Cas9/952₁;952₂ | 0.930 | 0.710 | < 0.0001 | *** |
| β2Cas9/952₂ | β2Cas9/739 | 0.930 | 0.857 | 0.3118 | ns |
| β2Cas9/952₁;952₂ | β2Cas9/739 | 0.710 | 0.857 | 0.0651 | ns |

Compact letter display (groups sharing a letter are not significantly different):

| Group | Mean | Letter(s) |
| --- | --- | --- |
| β2Cas9/952₁;952₂ | 0.710 | b |
| β2Cas9/739 | 0.857 | ab |
| WT/WT | 0.883 | ab |
| β2Cas9/952₂ | 0.930 | a |
| β2Cas9 | 0.962 | a |

#### Figure 2B - Pupa-to-adult survival

| Group | n (replicates) | Mean | SD |
| --- | --- | --- | --- |
| WT/WT | 3 | 0.823 | 0.115 |
| β2Cas9 | 3 | 0.943 | 0.012 |
| β2Cas9/952₂ | 5 | 0.936 | 0.040 |
| β2Cas9/952₁;952₂ | 4 | 0.887 | 0.025 |
| β2Cas9/739 | 3 | 0.920 | 0.010 |

Levene's test: F(4,13) = 3.3263, p = 0.0439 *

| Metric | Test | F | df | p |
| --- | --- | --- | --- | --- |
| Pupa-to-adult survival (Figure 2B) | Welch's one-way ANOVA | 3.7553 | (4, 6.02) | 0.0728 ns |

Overall test not significant; no post-hoc comparisons performed.

### Figure 4

Five genotypes: WT/WT, β2Cas9, zpgCas9, β2Cas9/wupA, zpgCas9/wupA. n = 3 replicates for WT/WT, β2Cas9, zpgCas9; n = 7 replicates for β2Cas9/wupA; n = 7 replicates for zpgCas9/wupA. Test selection based on Levene’s test.

### Figure 4A - % male pupae

| Group | n (replicates) | Mean | SD |
| --- | --- | --- | --- |
| WT/WT | 3 | 0.490 | 0.010 |
| β2Cas9 | 3 | 0.527 | 0.015 |
| zpgCas9 | 3 | 0.480 | 0.056 |
| β2Cas9/wupA | 7 | 0.983 | 0.017 |
| zpgCas9/wupA | 7 | 0.677 | 0.146 |

Levene's test: F(4,18) = 1.9511, p = 0.1456 ns

| Metric | Test | F | df | p |
| --- | --- | --- | --- | --- |
| % male pupae (Figure 4A) | One-way ANOVA | 30.7820 | (4, 18) | < 0.0001 *** |

Tukey HSD post-hoc pairwise comparisons:

| Group 1 | Group 2 | Mean 1 | Mean 2 | p | Sig. |
| --- | --- | --- | --- | --- | --- |
| WT/WT | β2Cas9 | 0.490 | 0.527 | 0.9846 | ns |
| WT/WT | zpgCas9 | 0.490 | 0.480 | 0.9999 | ns |
| WT/WT | β2Cas9/wupA | 0.490 | 0.983 | < 0.0001 | *** |
| WT/WT | zpgCas9/wupA | 0.490 | 0.677 | 0.0414 | * |
| β2Cas9 | zpgCas9 | 0.527 | 0.480 | 0.9630 | ns |
| β2Cas9 | β2Cas9/wupA | 0.527 | 0.983 | < 0.0001 | *** |
| β2Cas9 | zpgCas9/wupA | 0.527 | 0.677 | 0.1329 | ns |
| zpgCas9 | β2Cas9/wupA | 0.480 | 0.983 | < 0.0001 | *** |
| zpgCas9 | zpgCas9/wupA | 0.480 | 0.677 | 0.0295 | * |
| β2Cas9/wupA | zpgCas9/wupA | 0.983 | 0.677 | < 0.0001 | *** |

Compact letter display (groups sharing a letter are not significantly different):

| Group | Mean | Letter(s) |
| --- | --- | --- |
| zpgCas9 | 0.480 | a |
| WT/WT | 0.490 | a |
| β2Cas9 | 0.527 | ab |
| zpgCas9/wupA | 0.677 | b |
| β2Cas9/wupA | 0.983 | c |

### Figure 4A - % male adults

| Group | n (replicates) | Mean | SD |
| --- | --- | --- | --- |
| WT/WT | 3 | 0.467 | 0.035 |
| β2Cas9 | 3 | 0.520 | 0.010 |
| zpgCas9 | 3 | 0.480 | 0.075 |
| β2Cas9/wupA | 7 | 0.987 | 0.014 |
| zpgCas9/wupA | 7 | 0.886 | 0.113 |

Levene's test: F(4,18) = 6.3008, p = 0.0024 **

| Metric | Test | F | df | p |
| --- | --- | --- | --- | --- |
| % male adults (Figure 4A) | Welch's one-way ANOVA | 728.98 | (4, 6.04) | < 0.0001 *** |

Games–Howell post-hoc pairwise comparisons:

| Group 1 | Group 2 | Mean 1 | Mean 2 | p | Sig. |
| --- | --- | --- | --- | --- | --- |
| WT/WT | β2Cas9 | 0.467 | 0.520 | 0.3244 | ns |
| WT/WT | zpgCas9 | 0.467 | 0.480 | 0.9979 | ns |
| WT/WT | β2Cas9/wupA | 0.467 | 0.987 | 0.0027 | ** |
| WT/WT | zpgCas9/wupA | 0.467 | 0.886 | 0.0002 | *** |
| β2Cas9 | zpgCas9 | 0.520 | 0.480 | 0.8745 | ns |
| β2Cas9 | β2Cas9/wupA | 0.520 | 0.987 | < 0.0001 | *** |
| β2Cas9 | zpgCas9/wupA | 0.520 | 0.886 | 0.0007 | *** |
| zpgCas9 | β2Cas9/wupA | 0.480 | 0.987 | 0.0208 | * |
| zpgCas9 | zpgCas9/wupA | 0.480 | 0.886 | 0.0035 | ** |
| β2Cas9/wupA | zpgCas9/wupA | 0.987 | 0.886 | 0.2411 | ns |

Compact letter display (groups sharing a letter are not significantly different):

| Group | Mean | Letter(s) |
| --- | --- | --- |
| WT/WT | 0.467 | a |
| zpgCas9 | 0.480 | a |
| β2Cas9 | 0.520 | a |
| zpgCas9/wupA | 0.886 | b |
| β2Cas9/wupA | 0.987 | b |

### Figure 4B - Hatching rates

| Group | n (replicates) | Mean | SD |
| --- | --- | --- | --- |
| WT/WT | 3 | 0.913 | 0.091 |
| β2Cas9 | 3 | 0.907 | 0.025 |
| zpgCas9 | 3 | 0.937 | 0.078 |
| β2Cas9/wupA | 7 | 0.893 | 0.119 |
| zpgCas9/wupA | 7 | 0.759 | 0.201 |

Levene's test: F(4,18) = 1.5127, p = 0.2405 ns

| Metric | Test | F | df | p |
| --- | --- | --- | --- | --- |
| Hatching rate (Figure 4B) | One-way ANOVA | 1.4177 | (4, 18) | 0.2684 ns |

Overall test not significant; no post-hoc comparisons performed.

**Figure 4B - Larva-to-pupa survival**

| Group | n (replicates) | Mean | SD |
| --- | --- | --- | --- |
| WT/WT | 3 | 0.883 | 0.111 |
| β2Cas9 | 3 | 0.962 | 0.001 |
| zpgCas9 | 3 | 0.923 | 0.025 |
| β2Cas9/wupA | 7 | 0.900 | 0.079 |
| zpgCas9/wupA | 7 | 0.649 | 0.150 |

Levene's test: F(4,18) = 1.5112, p = 0.2409 ns

| Metric | Test | F | df | p |
| --- | --- | --- | --- | --- |
| Larva-to-pupa survival (Figure 4B) | One-way ANOVA | 8.0016 | (4, 18) | 0.0007 *** |

Tukey HSD post-hoc pairwise comparisons:

| Group 1 | Group 2 | Mean 1 | Mean 2 | p | Sig. |
| --- | --- | --- | --- | --- | --- |
| WT/WT | β2Cas9 | 0.883 | 0.962 | 0.8845 | ns |
| WT/WT | zpgCas9 | 0.883 | 0.923 | 0.9894 | ns |
| WT/WT | β2Cas9/wupA | 0.883 | 0.900 | 0.9993 | ns |
| WT/WT | zpgCas9/wupA | 0.883 | 0.649 | 0.0320 | * |
| β2Cas9 | zpgCas9 | 0.962 | 0.923 | 0.9904 | ns |
| β2Cas9 | β2Cas9/wupA | 0.962 | 0.900 | 0.9073 | ns |
| β2Cas9 | zpgCas9/wupA | 0.962 | 0.649 | 0.0032 | ** |
| zpgCas9 | β2Cas9/wupA | 0.923 | 0.900 | 0.9974 | ns |
| zpgCas9 | zpgCas9/wupA | 0.923 | 0.649 | 0.0101 | * |
| β2Cas9/wupA | zpgCas9/wupA | 0.900 | 0.649 | 0.0023 | ** |

Compact letter display (groups sharing a letter are not significantly different):

| Group | Mean | Letter(s) |
| --- | --- | --- |
| zpgCas9/wupA | 0.649 | b |
| WT/WT | 0.883 | a |
| β2Cas9/wupA | 0.900 | a |
| zpgCas9 | 0.923 | a |
| β2Cas9 | 0.962 | a |

### Figure 4B - Pupa-to-adult survival

| Group | n (replicates) | Mean | SD |
| --- | --- | --- | --- |
| WT/WT | 3 | 0.823 | 0.115 |
| β2Cas9 | 3 | 0.943 | 0.012 |
| zpgCas9 | 3 | 0.977 | 0.021 |
| β2Cas9/wupA | 7 | 0.901 | 0.036 |
| zpgCas9/wupA | 7 | 0.693 | 0.190 |

Levene's test: F(4,18) = 4.1915, p = 0.0143 *

| Metric | Test | F | df | p |
| --- | --- | --- | --- | --- |
| Pupa-to-adult survival (Figure 4B) | Welch's one-way ANOVA | 6.2970 | (4, 6.86) | 0.0187 * |

Games–Howell post-hoc pairwise comparisons:

| Group 1 | Group 2 | Mean 1 | Mean 2 | p | Sig. |
| --- | --- | --- | --- | --- | --- |
| WT/WT | β2Cas9 | 0.823 | 0.943 | 0.5361 | ns |
| WT/WT | zpgCas9 | 0.823 | 0.977 | 0.3953 | ns |
| WT/WT | β2Cas9/wupA | 0.823 | 0.901 | 0.7805 | ns |
| WT/WT | zpgCas9/wupA | 0.823 | 0.693 | 0.6833 | ns |
| β2Cas9 | zpgCas9 | 0.943 | 0.977 | 0.3038 | ns |
| β2Cas9 | β2Cas9/wupA | 0.943 | 0.901 | 0.1271 | ns |
| β2Cas9 | zpgCas9/wupA | 0.943 | 0.693 | 0.0672 | ns |
| zpgCas9 | β2Cas9/wupA | 0.977 | 0.901 | 0.0267 | * |
| zpgCas9 | zpgCas9/wupA | 0.977 | 0.693 | 0.0396 | * |
| β2Cas9/wupA | zpgCas9/wupA | 0.901 | 0.693 | 0.1326 | ns |

Compact letter display (groups sharing a letter are not significantly different):

| Group | Mean | Letter(s) |
| --- | --- | --- |
| zpgCas9/wupA | 0.693 | b |
| WT/WT | 0.823 | ab |
| β2Cas9/wupA | 0.901 | b |
| β2Cas9 | 0.943 | ab |
| zpgCas9 | 0.977 | a |

### Figure 6

Two-group comparisons between Y^RFP^/zpgCas9/wupA (experimental) and Y^RFP^/zpgCas9 (control). n = 3 replicates per group. All pairwise tests are two-tailed Welch’s t-tests.

### Figure 6B and 6C Sex ratio, hatching, and overall survival

| Comparison | Mean (Exp) | Mean (Ctrl) | t | df | p |
| --- | --- | --- | --- | --- | --- |
| Hatching rate | 0.900 | 0.767 | 4.126 | 3.58 | 0.0182 * |
| % male at L1 (Figure 6B) | 0.547 | 0.523 | 1.565 | 2.94 | 0.2172 ns |
| % male at pupae (Figure 6B) | 0.783 | 0.527 | 4.170 | 2.65 | 0.0318 * |
| % male at adults (Figure 6B) | 0.980 | 0.520 | 17.816 | 2.21 | 0.0020 ** |
| Overall L-P survival (Figure 6C) | 0.497 | 0.880 | 8.214 | 3.20 | 0.0030 ** |
| Overall P-A survival (Figure 6C) | 0.677 | 0.910 | 3.865 | 2.27 | 0.0494 * |

#### Figure 6D Sex-specific mortality (4-group comparison)

Four groups: Exp sons, Exp daughters, Ctrl sons, Ctrl daughters. Test selection (one-way ANOVA vs Welch’s) based on Levene’s test per stage.

##### Figure 6D - Larva-to-pupa mortality

| Group | n (replicates) | Mean | SD |
| --- | --- | --- | --- |
| Exp sons | 3 | 0.283 | 0.147 |
| Exp daughters | 3 | 0.767 | 0.095 |
| Ctrl sons | 3 | 0.113 | 0.090 |
| Ctrl daughters | 3 | 0.137 | 0.061 |

Levene's test: F(3,8) = 1.2767, p = 0.3464 ns

| Metric | Test | F | df | p |
| --- | --- | --- | --- | --- |
| Larva-to-pupa mortality | One-way ANOVA | 25.4868 | (3, 8) | 0.0002 *** |

Tukey HSD post-hoc pairwise comparisons:

| Group 1 | Group 2 | Mean 1 | Mean 2 | p | Sig. |
| --- | --- | --- | --- | --- | --- |
| Exp sons | Exp daughters | 0.283 | 0.767 | 0.0020 | ** |
| Exp sons | Ctrl sons | 0.283 | 0.113 | 0.2725 | ns |
| Exp sons | Ctrl daughters | 0.283 | 0.137 | 0.3824 | ns |
| Exp daughters | Ctrl sons | 0.767 | 0.113 | 0.0003 | *** |
| Exp daughters | Ctrl daughters | 0.767 | 0.137 | 0.0003 | *** |
| Ctrl sons | Ctrl daughters | 0.113 | 0.137 | 0.9922 | ns |

Compact letter display (groups sharing a letter are not significantly different):

| Group | Mean | Letter(s) |
| --- | --- | --- |
| Ctrl sons | 0.113 | a |
| Ctrl daughters | 0.137 | a |
| Exp sons | 0.283 | a |
| Exp daughters | 0.767 | b |

### Figure 6D Pupa-to-adult mortality

| Group | n (replicates) | Mean | SD |
| --- | --- | --- | --- |
| Exp sons | 3 | 0.157 | 0.032 |
| Exp daughters | 3 | 0.920 | 0.078 |
| Ctrl sons | 3 | 0.100 | 0.026 |
| Ctrl daughters | 3 | 0.083 | 0.035 |

Levene's test: F(3,8) = 3.1754, p = 0.0850 ns

| Metric | Test | F | df | p |
| --- | --- | --- | --- | --- |
| Pupa-to-adult mortality | One-way ANOVA | 234.3913 | (3, 8) | < 0.0001 *** |

Tukey HSD post-hoc pairwise comparisons:

| Group 1 | Group 2 | Mean 1 | Mean 2 | p | Sig. |
| --- | --- | --- | --- | --- | --- |
| Exp sons | Exp daughters | 0.157 | 0.920 | < 0.0001 | *** |
| Exp sons | Ctrl sons | 0.157 | 0.100 | 0.4761 | ns |
| Exp sons | Ctrl daughters | 0.157 | 0.083 | 0.2802 | ns |
| Exp daughters | Ctrl sons | 0.920 | 0.100 | < 0.0001 | *** |
| Exp daughters | Ctrl daughters | 0.920 | 0.083 | < 0.0001 | *** |
| Ctrl sons | Ctrl daughters | 0.100 | 0.083 | 0.9685 | ns |

Compact letter display (groups sharing a letter are not significantly different):

| Group | Mean | Letter(s) |
| --- | --- | --- |
| Ctrl daughters | 0.083 | a |
| Ctrl sons | 0.100 | a |
| Exp sons | 0.157 | a |
| Exp daughters | 0.920 | b |

#### Figure 6F - Mean day of pupation

Four groups: Ctrl sons, Ctrl daughters, Exp sons, Exp daughters. Pairwise Welch’s t-tests.

| Group | n (replicates) | Mean | SD |
| --- | --- | --- | --- |
| Ctrl sons | 3 | 7.880 | 0.874 |
| Ctrl daughters | 3 | 8.166 | 0.481 |
| Exp sons | 3 | 9.122 | 0.047 |
| Exp daughters | 3 | 10.260 | 0.457 |

| Comparison | Mean 1 (days) | Mean 2 (days) | t | df | p |
| --- | --- | --- | --- | --- | --- |
| Ctrl sons vs Ctrl daughters | 7.88 | 8.17 | -0.497 | 3.11 | 0.6521 ns |
| Exp sons vs Ctrl sons | 9.12 | 7.88 | 2.458 | 2.01 | 0.1325 ns |
| Exp daughters vs Ctrl daughters | 10.26 | 8.17 | 5.462 | 3.99 | 0.0055 ** |
| Exp daughters vs Exp sons | 10.26 | 9.12 | 4.285 | 2.04 | 0.0485 * |
